# Loss of Secretory Pathway Ca^2+^ ATPase (SPCA1) Impairs Insulin Secretion and Reduces Autophagy in the Pancreatic Islet

**DOI:** 10.1101/2021.08.30.458203

**Authors:** Robert N. Bone, Xin Tong, Staci A. Weaver, Charanya Muralidharan, Preethi Krishnan, Tatsuyoshi Kono, Carmella Evans-Molina

**Author notes:** Corresponding Author: Carmella Evans-Molina, MD, PhD, Indiana University School of Medicine, 635 Barnhill Drive MS 2031A, Indianapolis, IN, USA.

## Abstract

The β cell Golgi apparatus serves as a significant store of intracellular Ca^2+^ and an important site of proinsulin maturation. However, the contribution of Golgi Ca^2+^ to diabetes pathophysiology is unknown. The Golgi primarily utilizes the Secretory Pathway Ca^2+^ ATPase (SPCA1) to maintain intraluminal Ca^2+^ stores, and loss of SPCA1 has been linked to impaired Golgi function in other cell types. Here, we demonstrated that SPCA1 expression is decreased in islets from diabetic mice and human organ donors with type 2 diabetes, suggesting SPCA1 may impact diabetes development. INS-1 β cells lacking SPCA1 (SPCA1KO) showed reduced intraluminal Golgi Ca^2+^ levels, reduced glucose-stimulated insulin secretion (GSIS), and increased insulin content. Islets from SPCA1 haploinsufficient mice (SPCA1^+/-^) exhibited reduced GSIS, altered glucose-induced Ca^2+^ oscillations, and altered insulin granule maturation. Autophagy can regulate granule homeostasis, therefore we induced autophagy with Torin1 and found that SPCA1KO cells and SPCA1^+/-^ islets had reduced levels of the autophagosome marker LC3-II. Furthermore, SPCA1KO LC3-II were unchanged after blocking autophagy initiation or autophagolysosome fusion and acidification. Thus, we concluded that β cell SPCA1 plays an important role in the maintenance of Golgi Ca^2+^ homeostasis and reduced Golgi Ca^2+^ impairs autophagy initiation and may impact insulin granule homeostasis.

## INTRODUCTION

In order to regulate minute-to-minute fluctuations in blood glucose levels, the pancreatic β cell relies upon a highly developed secretory pathway capable of producing up to one million insulin molecules per minute [1]. Insulin production begins within the lumen of the endoplasmic reticulum (ER), where signal peptidase cleaves the preproinsulin signal sequence to convert preproinsulin to proinsulin. Subsequently, proinsulin is trafficked out of the ER to the Golgi apparatus, where it is packaged into early secretory vesicles [2]. Cleavage of proinsulin by prohormone convertases to produce mature insulin begins within the trans-Golgi sub-compartment and is completed within the secretory vesicles [2]. The organelles of the secretory pathway contain high intraluminal levels of Ca^2+^: 300-700 µM in the ER, ∼300 µM in the Golgi apparatus, and ∼50 µM in the secretory vesicles [3, 4]. In contrast, cytosolic Ca^2+^ concentration is at least three orders of magnitude lower. These steep concentration gradients are primarily maintained by two P-type ATPases: the sarco/endoplasmic reticulum Ca^2+^ ATPase (SERCA) in the ER and cis-Golgi and the secretory pathway Ca^2+^ ATPase (SPCA) in the Golgi apparatus. We and others have previously identified loss of SERCA2 expression and activity in models of type 1 and type 2 diabetes and have linked SERCA2 loss with glucose intolerance, reduced insulin secretion, decreased β cell proliferation, and increased β cell ER stress [5-7]. Within the Golgi, Ca^2+^ controls several key steps relevant to insulin production and organelle homeostasis, including protein trafficking, cargo condensation, protein secretion, and enzymatic processing of proteins, such as the conversion of proinsulin to insulin by prohormone convertases [8, 9]. However, to date, the contribution of Golgi apparatus Ca^2+^ and SPCA to Ca^2+^ homeostasis in the secretory pathway of β cells remains largely unknown.

The SPCA pump transports one Ca^2+^ or one Mn^2+^ per one ATP hydrolyzed, with a much higher affinity for Ca^2+^ than SERCA [10, 11]. Two isoforms of SPCA have been identified, SPCA1 and SPCA2, that are encoded by *ATP2C1* and *ATP2C2* genes, respectively [12]. SPCA1 is ubiquitously expressed with higher expression found in secretory cells of the salivary, adrenal, and mammary glands, testis, and the skin [13, 14]. In contrast, the distribution of SPCA2 is more variable and known to be expressed in many types of secretory cells [14]. Both SPCA1 and SPCA2 are expressed in human islets, with SPCA1 at a higher level (TIGER Data Portal - http://tiger.bsc.es; [15]) Within the secretory pathway, SPCA1 expression is localized to the cis-, medial-, and trans-Golgi, with increasing concentration towards the trans-Golgi, and to the secretory vesicles [11, 16].

Loss of SPCA1 and reduced concentration of Ca^2+^ in the Golgi apparatus has been associated with impaired Golgi functions, including glycosylation, proteolytic processing, and cargo trafficking, and increased apoptosis in several cell types [12]. Thus far, only one study has addressed the role of SPCA1 in the β cell, demonstrating that the acute knockdown of SPCA1 in MIN6 and INS-1 β cell lines leads to reduced Ca^2+^ oscillations and increased glucose-induced insulin secretion; however, the mechanisms underlying these changes were not identified [17]. Given these gaps in understanding of how alterations in Golgi Ca^2+^ and loss of SPCA1 may impact β cell function and Ca^2+^ homeostasis, we have generated a SPCA1-knockout INS-1 cell line and a whole-body SPCA1-haploinsufficient mouse model to test the hypothesis that disruptions in Golgi Ca^2+^ homeostasis may impact insulin granule homeostasis leading to impaired β cell function.

## RESEARCH DESIGN AND METHODS

### Animal and Islet Studies

SPCA1^+/-^ (MMRRC stock #36808 [18]), Akita, db/db, and C57BL/6J mice were obtained from Jackson Laboratory (Bar Harbor, ME). SPCA^+/-^ mice were backcrossed onto the C57BL/6J background for a minimum of ten generations prior to use. Mice were maintained under protocols approved by the Indiana University IACUC. Cages were kept in a standard light-dark cycle with *ad libitum* access to food and water.

Lean and fat mass were estimated using an EchoMRI Body Composition Analyzer (EchoMRI LLC, Houston, TX). Intraperitoneal glucose tolerance tests (GTT) were performed after 16 h of fasting, using a glucose dose of 2 g/kg lean body mass. Insulin tolerance tests (ITT) were performed after 6 h of fasting, using regular human insulin (NovoNordisk, Plainsboro, NJ) at a dose of 0.75 U/kg lean body mass. Blood glucose levels were measured using the AlphaTRAK glucometer (Abbott Laboratories, Abbott Park, IL).

Mouse islets were isolated by collagenase digestion as previously described [19]. Glucose-stimulated insulin secretion (GSIS) in isolated islets was measured using the Biorep Perifusion System (Biorep, Miami Lakes, FL) [20]. Handpicked islets (50/chamber) were perifused with Krebs buffer containing 2.8 mM glucose for 20 min, followed by 16.7 mM glucose for 30 min and 30 mM KCl for 30 min at a rate of 120 μL/min. Secreted insulin was measured by ELISA (Mercodia, Uppsala, Sweden), and results were normalized to DNA content, measured using the Quant-iT PicoGreen dsDNA Assay Kit (ThermoFisher, Waltham, MA).

For ultrastructural analysis, isolated islets were fixed and transferred to the Advanced Electron Microscopy Facility at the University of Chicago. Insulin granule maturity states were quantitated manually in transmission electron microscopy images using ImageJ software (NIH, Bethesda, MD) as previously described [21].

Human cadaveric donor islets were obtained from the Integrated Islet Distribution Program (Supporting Information Table S1) [22]. Upon receipt, islets were allowed to recover overnight in Islet Media. Islets were collected, washed in PBS, and either lysed in RLT+ buffer (Qiagen, Hilden, Germany) prior to total RNA isolation or lysed in protein lysis buffer prior to immunoblotting. Further information can be found in the Supporting Information.

### Cell Culture

Rat INS-1 832/13 cells (WT) were cultured as previously described [23]. CRISPR/Cas9 genomic editing was used to create a SPCA1 knockout INS-1 832/13 cell line (SPCA1KO) at the Genome Engineering and iPSC Center at Washington University in St. Louis. Guide RNAs (AGTCATAGGCGAGCCTTCCANGG and ATAGGCGAGCCTTCCATGGCNGG) were used to disable *Atp2c1*.

For GSIS, INS-1 cells were seeded in 12-well plates and allowed to reach 80-90% confluency. Cells were preincubated in Krebs buffer containing 2.5 mM glucose for 2 h and then switched to Krebs buffer containing either 2.5 mM or 15 mM glucose for 2 h. Supernatants were collected, and insulin concentration was measured by radioimmunoassay (RIA) (MilliporeSigma, Burlington, MA) and total cellular insulin content was normalized to total protein.

### Immunoblotting, Reverse Phase Protein Array, qRT-PCR, Immunofluorescence, and β cell area

Immunoblotting was performed as previously described [24, 25]. In brief, cultured cells or isolated islets were washed with PBS and lysed with protein lysis buffer. Lysates (20 μg) were separated by SDS-PAGE, transferred to a PVDF membrane, and incubated overnight at 4°C with primary antibodies (Supporting Information Table S2). Bound primary antibodies were detected with IRDye-conjugated secondary antibodies and visualized using fluorometric scanning on an Odyssey Imaging System (LI-COR Biosciences, Lincoln, NE).

For the Reverse Phase Protein Array (RPPA), cells were washed in PBS and lysed in RPPA lysis buffer. Lysates were adjusted to 1.5 μg protein/mL and denatured with RPPA 4x sample buffer. Samples were transferred to the Functional Proteomics RPPA Core Facility at The University of Texas MD Anderson Cancer Center (Houston, TX). Biological pathways were identified using Metascape 3.5 (metascape.org) [26], and pathways with *P* < 0.05 were considered significant.

Quantitative real-time (qRT)-PCR was performed using primers listed in Supporting Information Table S3. In brief, total RNA was extracted from cultured cells or isolated islets using RNeasy Plus Mini or Micro kits (Qiagen, Valencia, CA) according to the manufacturer’s instructions. cDNA was generated via reverse transcription for 1 h at 37°C using random hexamers, 0.5 mM dNTPs, 5x first-strand buffer, 0.01 mM DTT, and MML-V reverse transcriptase (all from Invitrogen, Carlsbad, CA). qRT-PCR reactions were performed, and relative RNA levels were calculated using the comparative C_T_ method normalized to *Actb*, as previously described [24, 25].

Immunofluorescently labeled slides were visualized with a Zeiss confocal microscope, and z-stack images were acquired. For Golgi volumetric analysis, z-stack images were analyzed using ImageJ. For LC3 puncta and colocalization analysis, CellProfiler 4.1.3 (cellprofiler.org) [27] was used as previously described [28]. Further information can be found in the Supporting Information.

For β cell area, pancreases were processed, and immunohistochemistry was performed as previously described [29] and further information can be found in the Supporting Information. Sections, 3-5 per mouse and at least 50 μm apart, were analyzed using Zen Blue software (Zeiss, Oberkochen, Germany). The insulin-positive area in pixels was divided by the total pancreas area in pixels and expressed as a percentage.

### Live Cell Imaging

Cytosolic Ca^2+^ dynamics in WT and SPCA1KO INS-1 cells were measured using the FLIPR Calcium 6 Assay Kit and a FlexStation 3 plate reader (Molecular Devices, Sunnyvale, CA) as previously described [30]. In brief, two days after cell seeding, Calcium 6 reagent was added and incubated for 2 h at 37°C. After washing with modified HBSS (see Supporting Information), fluorescence was recorded before and after drug addition. The ΔF/F0 was then calculated from the resting intercellular Ca^2+^ (F0) and drug response (ΔF).

Isolated islet cytosolic Ca^2+^ dynamics were measured using the ratiometric Ca^2+^ indicator fura-2-acetoxymethyl ester (fura2-AM) and imaged using a Zeiss Z1 microscope, as previously described [25]. In brief, isolated islets were incubated in 2.5 mM glucose Ca^2+^ Imaging HBSS with 5 μM fura2-AM for 30 min at 37°C/5% CO_2_. Islets were transferred to a glass-bottom dish (MatTek Corp., Ashland, MA), and baseline fura2-AM fluorescence ratio (340/380 nm) was recorded. High glucose-supplemented HBSS was added to reach a final concentration of 15 mM glucose, and the fura2-AM fluorescence ratio was recorded. Individual islet traces were graphed, and amplitudes (nadir-to-peak) and periods (nadir-to-nadir) were measured with ImageJ.

Intraluminal Golgi Ca^2+^ differences were determined in INS-1 cells transduced with a Golgi compartment-directed Ca^2+^ biosensor adenovirus, and the signal was monitored using fluorescence lifetime imaging microscopy (FLIM), as previously described [24]. Golgi compartment-specific adenoviruses expressing Cameleon calcium sensor probes specific for the medial-Golgi, driven by the 1,6 N acetyl-glucosaminyl transferase promoter [16] and the trans-Golgi, driven by the sialyl-transferase1 promoter [31], were generated by the Human Islet and Adenovirus Core at Einstein-Mount Sinai Diabetes Research Center (Bronx, NY). The cells were transduced with the corresponding adenovirus 24 h prior to imaging. Confocal images were acquired using an Alba FastFLIM system with VistaVision software (ISS Inc., Champaign, IL). Further information can be found in the Supporting Information.

### Statistical Analysis

The significance of differences between groups was determined using the unpaired Student *t*-test, one-way ANOVA, or two-way ANOVA. Results are reported as the mean ± SEM. GraphPad Prism software (GraphPad Software, Inc., La Jolla, CA) was used for data analysis. A *P-*value < 0.05 was considered to indicate a significant difference between groups.

## RESULTS

### SPCA1 Is Reduced in Models of Diabetes

To determine the predominant SPCA isoform expressed in the β cell, qRT-PCR was performed in human and mouse islets (Fig. 1*A-B*) and INS-1 cells (Fig. 1*C*) using validated primers for SPCA1 and SPCA2. In all three models, SPCA1 was detected at a significantly higher level than SPCA2. Next, we measured SPCA1 expression in islets from organ donors with type 2 diabetes (T2D), islets from Akita mice (a model of β cell ER stress), and islets from db/db mice (a model of obesity and hyperglycemia). Compared to islets from non-diabetic human cadaveric donors, SPCA1 mRNA (Fig. 1*D*) and protein (Fig. 1*E-F*) levels were reduced in islets from donors with T2D. Similarly, SPCA1 mRNA was reduced in islets from 6 wk old Akita mice (Fig. 1*G*) and SPCA1 protein was reduced in islets from 8 wk old db/db mice (Fig. 1*H-I*).

**Figure 1.**
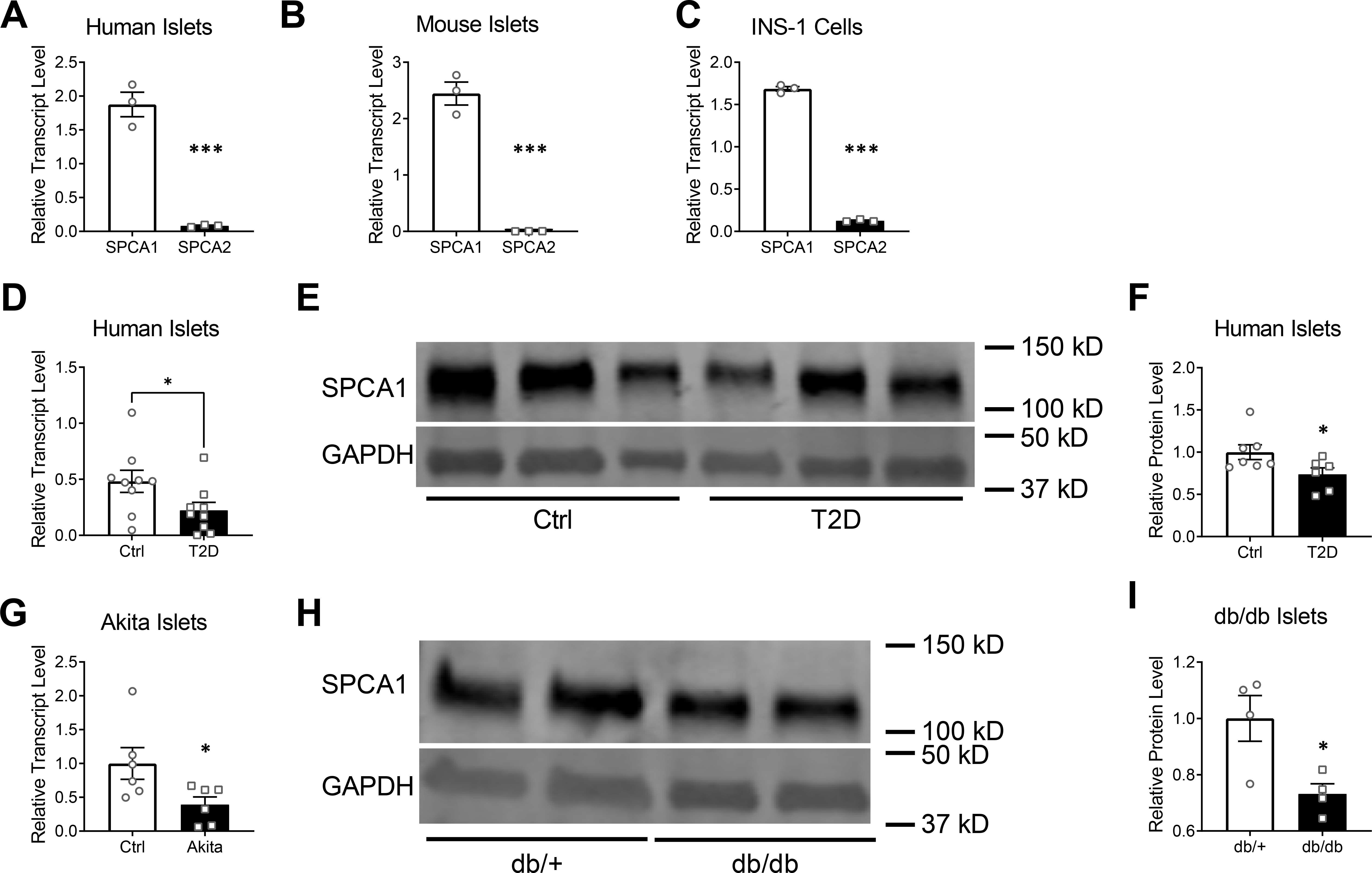
SPCA1 expression is reduced in models of diabetes. (*A*-*C*) qRT-PCR was performed to detect SPCA isoforms. SPCA1 is the predominant SPCA isoform in human islets (*A*), mouse islets (*B*), and INS-1 β cells (*C*); *n* = 3 per group, \*\*\**P* < 0.001, unpaired *t*-test. (*D-F*) SPCA1 expression in human islets from non-diabetic donors (Ctrl) and donors with type 2 diabetes (T2D) was determined by qRT-PCR normalized to actin (*D*) and Western blotting normalized to GAPDH (*E-F*); *n* = 9 per group (average age: Ctrl 44.78 ± 4.49 yr; T2D 53.22 ± 2.184 yr), \**P* < 0.05, unpaired *t*-test. (*G*) SPCA1 expression in mouse islets from 6 wk old control littermate and Akita mice was determined by qRT-PCR. *n* = 6 per group, \**P* < 0.05, unpaired *t*-test. (*H*-*I*) SPCA1 protein expression in islets from db/+ control littermate and db/db mice determined by Western blotting normalized to GAPDH; *n* = 4 per group, \**P* < 0.05, unpaired *t*-test.

### Golgi Ca^2+^ Concentration Is Reduced Following SPCA1 Loss

To determine how chronic SPCA1 deficiency impacts intraluminal Golgi Ca^2+^ regulation, we used CRISPR/Cas9 technology to generate INS-1 cells in which the SPCA1-encoding gene *Atp2c1* was disabled (SPCA1KO; Fig. 2*A*). To measure intra-Golgi Ca^2+^, we utilized highly specific Cameleon probes for the medial-Golgi and trans-Golgi sub-compartments. The donor probe lifetime was recorded using Fluorescence Lifetime Imaging Microscopy (FLIM). Higher Ca^2+^ concentration leads to increased binding to calmodulin within the Cameleon probe, causing enhanced emission and reduced donor lifetime. In SPCA1KO INS-1 cells, the fluorescence lifetime was higher in both sub-compartments, indicative of reduced Golgi Ca^2+^ with SPCA1 loss (Fig. 2*B-E*). To verify that the loss of Golgi Ca^2+^ was due to SPCA1 loss, we transduced SPCA1KO INS-1 cells with an adenovirus expressing human SPCA1 (Fig. 2*F*). As expected, restoration of SPCA1 normalized fluorescence lifetimes in the Golgi sub-compartments (Fig. 2*G-J*). To confirm that the decrease in Golgi Ca^2+^ was due to the loss of SPCA1 and not decreased Golgi volume, we used immunofluorescence to label markers of the Golgi apparatus and performed volumetric analysis using ImageJ. GM130 (Fig. 2*K*), giantin (Fig. 2*L*), and syntaxin6 (Fig. 2*M*) were used to label the cis-, medial-, and trans-Golgi, respectively [32, 33]. This analysis indicated that Golgi volume was not decreased in SPCA1KO INS-1 cells (Fig. 2*N-P*).

**Figure 2.**
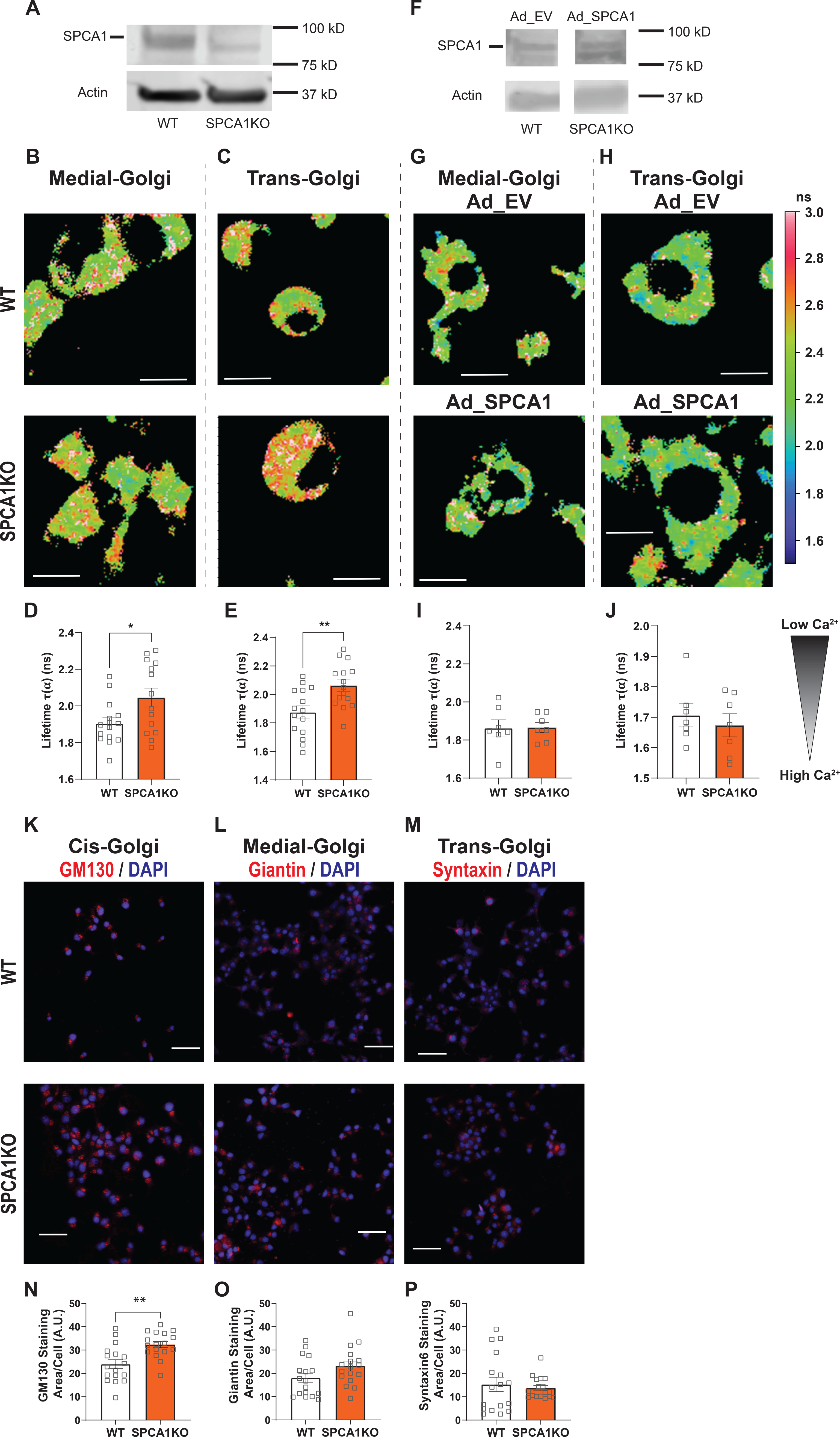
Loss of SPCA1 alters Ca^2+^ dynamics. (*A*) SPCA1 expression in WT and SPCA1KO INS-1 cells. (*B-E*) Fluorescent lifetime imaging microscopy (FLIM) in WT and SPCA1KO INS-1 cells. Cells were virally transduced with Golgi subcompartment-specific Cameleon donor probes (8 x 10^8^ viral particles/mL), and FLIM was used to measure medial-Golgi (*B*) and trans-Golgi (*C*) Ca^2+^ levels. Images show representative fluorescence lifetime maps with heatmap indicating donor lifetime in nanoseconds (ns). Scale bar = 10 μm. (*D-E*) Quantitation of fluorescence lifetime wherein higher probe lifetime corresponds to lower Ca^2+^ level; *n* = 14-15 fields of view containing at least 6 cells per field, * *P*<0.05, unpaired *t*-test. (*F***-***J*) Measurements of Ca^2+^ levels by FLIM in WT cells transfected with an empty adenoviral vector (Ad_EV; 2.5 x 10^6^ viral particles/mL) and SPCA1KO INS-1 cells transfected with an adenoviral SPCA1 vector (Ad_SPCA1; 2.5 x 10^6^ viral particles/mL) to restore SPCA1 (*F*). Representative fluorescence lifetime maps reflecting medial-Golgi (*G*) and trans-Golgi (*H*) Ca^2+^ levels. (*I-J*) Quantitation of fluorescence lifetime wherein higher probe lifetime corresponds to lower Ca^2+^ levels; *n* = 7 fields of view containing at least 5 cells per field. (*K-P*) Volumetric analysis of the Golgi apparatus. (*K-M*) Representative immunofluorescent images of cis-Golgi (*K*), medial-Golgi (*L*), and trans-Golgi (*M*). Scale bar = 20 μm. (*N-P*) Cellular content of cis-Golgi (*N*), medial-Golgi (*O*), and trans-Golgi (*P*); *n* = 5 per group, ** *P* < 0.01, unpaired *t*-test.

### Loss of SPCA1 Alters Intracellular Ca^2+^ Transport and β cell Function

Next, we investigated the functional impact of SPCA1 loss. Using the cytosolic Ca^2+^ dye Calcium 6, we measured baseline cytosolic Ca^2+^ levels and Ca^2+^ flux in response to selected stimuli. This analysis revealed that the baseline cytosolic Ca^2+^ concentration was higher in SPCA1KO INS-1 cells compared to wild-type (WT) INS-1 cells (Fig. 3*A*). Caffeine and carbachol were used to stimulate, respectively, RyR- and IP3R-mediated Ca^2+^ release from intracellular stores, and thapsigargin and cyclopiazonic acid (CPA) were each used to inhibit the SERCA2 pump. SPCA1KO INS-1 cells had reduced Ca^2+^ efflux in response to carbachol but not in response to caffeine, and reduced Ca^2+^ responses to thapsigargin and CPA (Fig. 3*B*).These findings supported the notion that secretory pathway intraluminal Ca^2+^ stores were reduced in SPCA1KO INS-1 cells. Next, static GSIS was measured and found to be reduced in SPCA1KO INS-1 cells compared to WT cells (Fig. 3*C*). Interestingly, insulin content in SPCA1KO INS-1 cells was increased at both the transcript (Fig. 3*D*) and protein (Fig. 3*E*) level.

**Figure 3.**
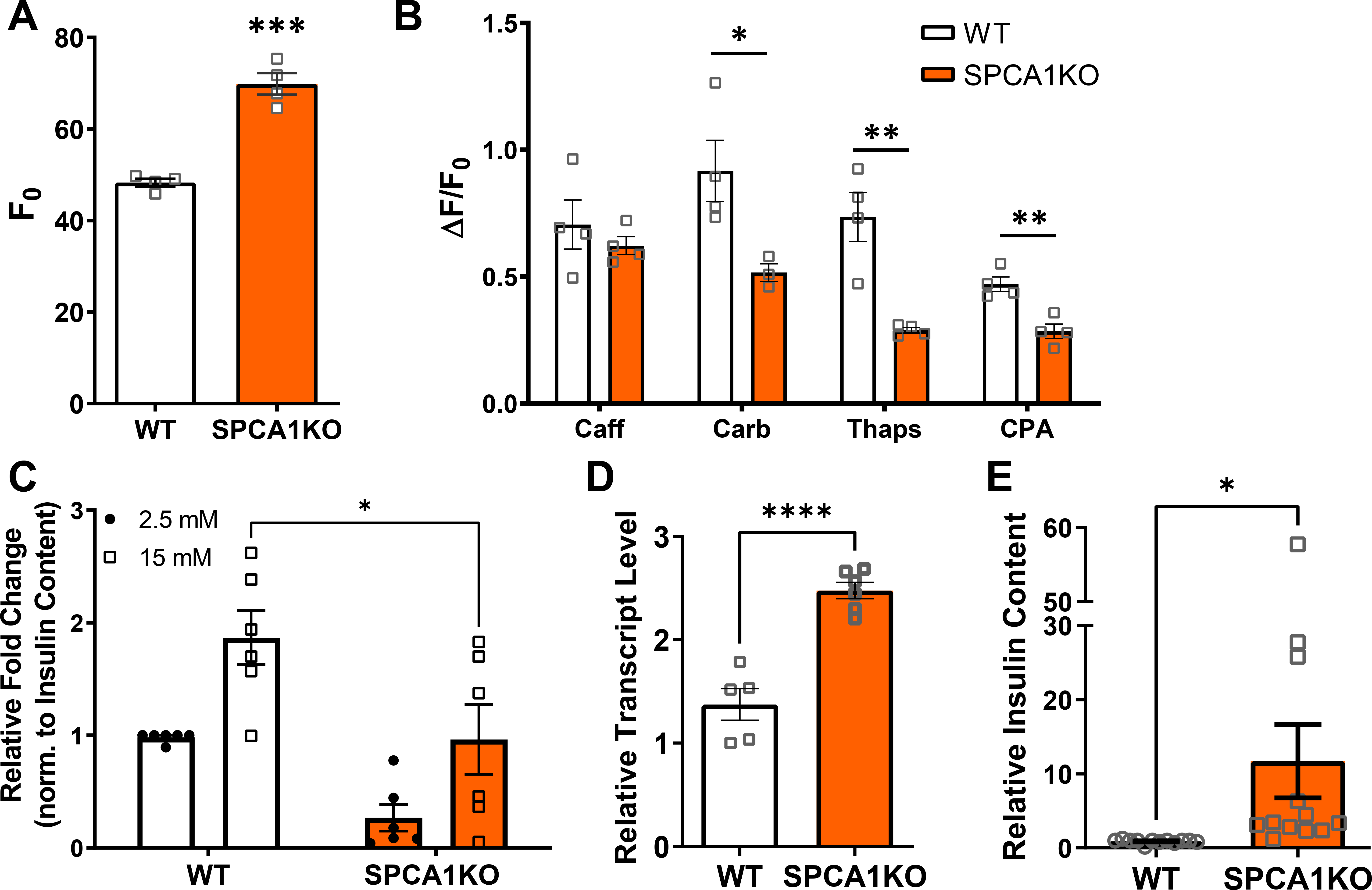
SPCA1 loss impairs insulin secretion. (*A*-*B*) Calcium imaging was performed in WT and SPCA1KO INS-1 cells using the cytosolic Ca^2+^ dye Calcium 6. (*A*) Average baseline (F_0_) cytosolic Ca^2+^; n = 4, \*\*\**P* < 0.001, unpaired *t*-test. (*B*) Change in Ca^2+^ (ΔF) in response to 5 mM caffeine (Caff), 100 μM carbachol (Carb), 10 μM thapsigargin (Thaps), and 0.1 μM cyclopiazonic acid (CPA), normalized to baseline cytosolic Ca^2+^ level (F_0_); *n* = 3-4, \**P* < 0.05, \*\**P* < 0.01, unpaired *t*-tests. (*C*) Glucose-stimulated insulin secretion in WT and SPCA1KO INS-1 cells. The concentration of secreted insulin was normalized to total cellular insulin content; *n* = 6, \**P* < 0.05, two-way ANOVA. (*D*) Insulin transcript level in WT and SPCA1KO INS-1 cells measured by qRT-PCR; *n* = 5-6, \*\*\*\**P* < 0.0001, unpaired *t*-test. (*E*) Insulin content in WT and SPCA1KO INS-1 cells measured by human insulin RIA, normalized to total protein; *n* = 12, \**P* < 0.05, unpaired *t*-test.

### Glucose Sensitivity and Body Composition are Unaffected by SPCA1 Haploinsufficiency

To interrogate the effects of SPCA1 deficiency in vivo, we utilized SPCA1 haploinsufficient mice (SPCA1^+/-^), since SPCA1^-/-^ mice do not survive past E10.5 due to the failure of the neural tube to close [18]. At 8 and 24 wks of age, SPCA1^+/-^ mice did not exhibit changes in body composition, glucose tolerance, or β cell mass compared to WT littermates (Fig. 4). SPCA1^+/-^ mice challenged with a high-fat diet (45% kcal from fat) for 16 wks also did not exhibit changes in these same parameters compared to WT littermates (Fig. S1).

**Figure 4.**
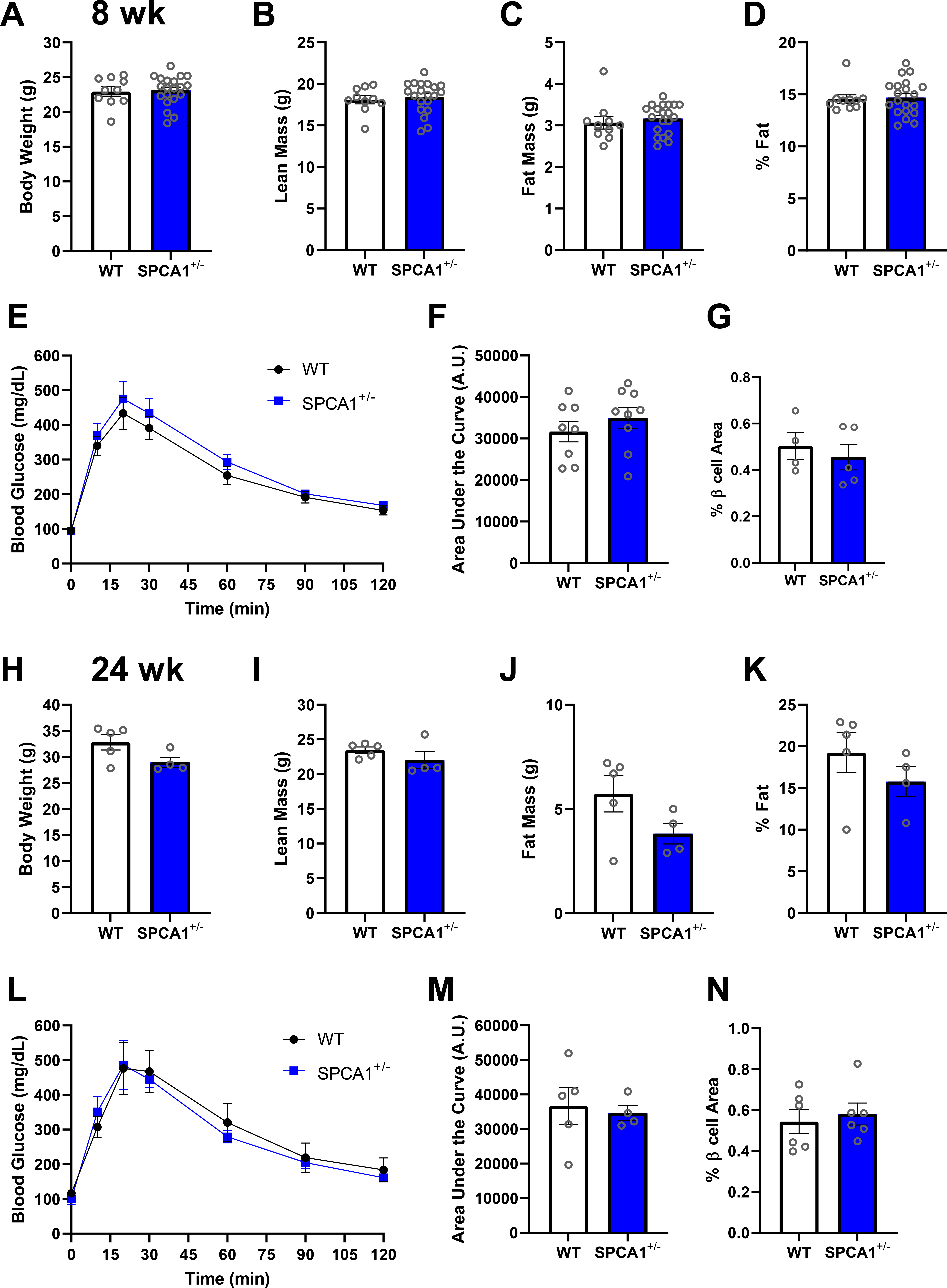
Body composition and glucose tolerance. (*A*-*D*) Body composition of 8 wk old male WT and SPCA1^+/-^ mice measured by EchoMRI. (*A*) Body weight; *n* = 10-20. (*B*) Lean mass; *n* = 10-20. (*C*) Fat mass; *n* = 10-20. (*D*) Percent fat; *n* = 10-20. (*E*-*F*) Glucose tolerance test in 8 wk old male WT and SPCA1^+/-^ mice (*E*) and corresponding area under the curve (*F*); *n* = 8-9. (*G*) Quantification of the fractional area of β cells in 8 wk old male WT and SPCA1^+/-^ mice; *n* = 4-5. (*H-K*) Body composition of 24 wk old male WT and SPCA1^+/-^ mice measured by EchoMRI. (*H*) Body weight; *n* = 4-5. (*I*) Lean mass; *n* = 4-5. (*J*) Fat mass; *n* = 4-5. (*K*) Percent fat; *n* = 4-5. (*L*-*M*) Glucose tolerance test of 24 wk old male WT and SPCA1^+/-^ mice (*L*) and corresponding area under the curve (*M*); *n* = 4-5. (*N*) Quantification of the fractional area of β cells in 24 wk old male WT and SPCA1^+/-^ mice; *n* = 6.

### Reduction in SPCA1-mediated Ca^2+^ Transport Impacts Insulin Secretion and Ca^2+^ Dynamics

Because calcium-induced Ca^2+^ release from secretory pathway stores is thought to amplify glucose-induced insulin exocytosis [34], we next investigated insulin secretion in isolated islets with reduced SPCA1. Despite no evidence of a whole-body phenotype, islets from SPCA1^+/-^ mice exhibited reduced first phase GSIS and reduced insulin release in response to KCl compared to islets from WT littermates (Fig. 5*A-D*).

**Figure 5.**
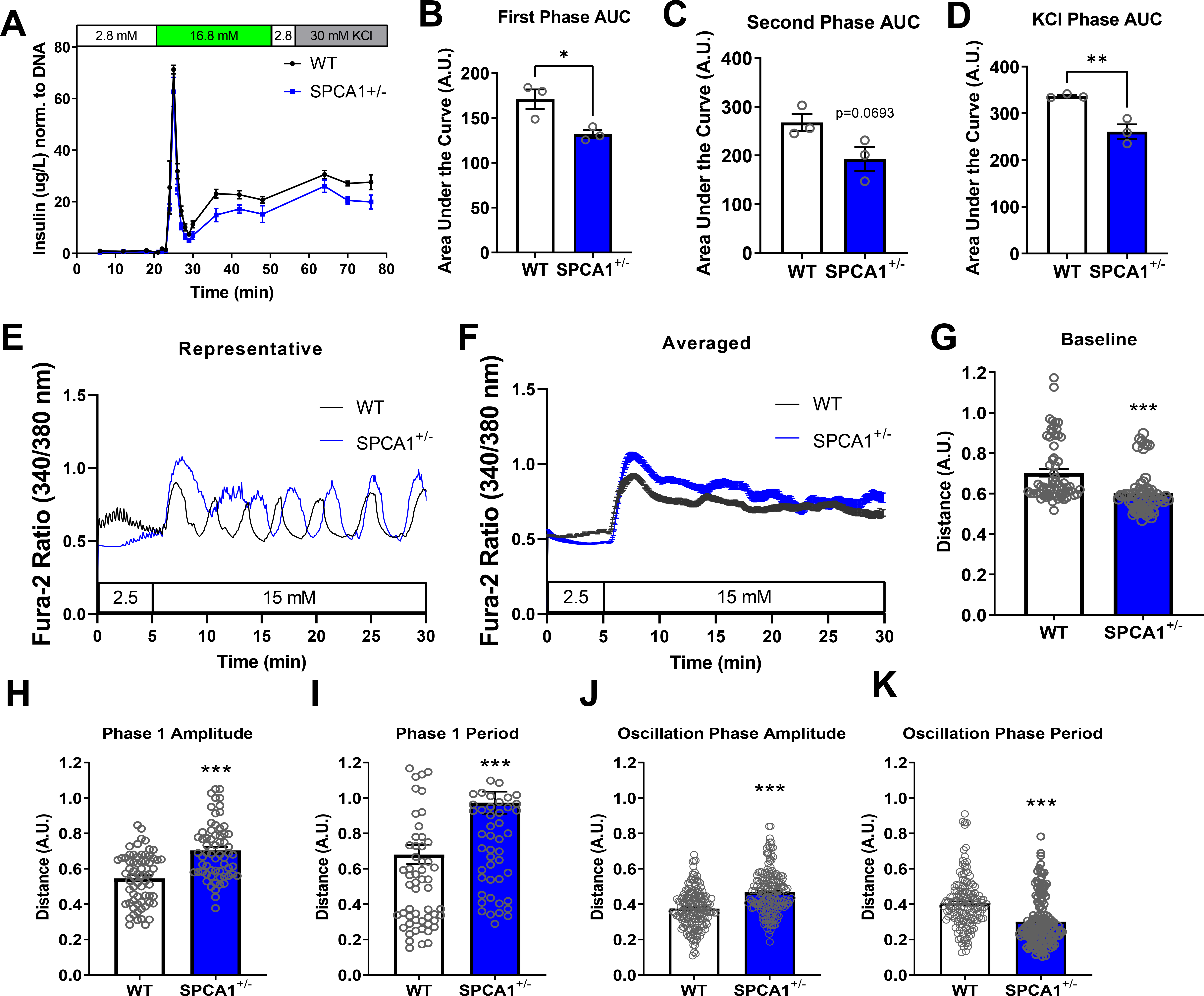
SPCA1 reduction impairs insulin secretion and alters calcium oscillations. (*A-D*) Islets isolated from 8 wk old WT and SPCA1^+/-^ mice were perifused with 2.8 mM glucose (minutes 0-20), 16.8 mM glucose (minutes 20-50), 2.8 mM glucose (minutes 50-56), and 30 mM KCl (minutes 56-76); *n* = 3. (*A*) Glucose-stimulated insulin secretion from isolated islets normalized to total islet DNA. (*B-D*) Area under the curve (AUC) analysis of insulin secretion during the first phase (*B*, minutes 20-30, \**P* < 0.05, unpaired *t*-test), second phase (*C*, minutes 30-50), and KCl phase (*D*, \*\**P* < 0.01, unpaired *t*-test). (*E-K*) Calcium oscillations in response to glucose were measured in islets isolated from 8 wk old male WT and SPCA1^+/-^ mice, using the ratiometric cytosolic calcium indicator fura2-AM. (*E*-*F*) Representative (*E*) and averaged (*F*) cytosolic Ca^2+^ recordings after stimulation with 15 mM glucose. (*G*) Average baseline cytosolic Ca^2+^. Average amplitude (*H*) and period (*I*) during the phase 1 response to glucose. Average amplitude (*J*) and period (*K*) during the oscillatory phase response to glucose. *E-K*, *n* = 63-66 isolated islets from 5 biological replicates per group; \*\*\**P* < 0.001, unpaired *t*-tests.

To test how these differences in insulin secretion correlated with Ca^2+^ signaling, the ratiometric cytosolic calcium indicator fura-2AM was used (Fig. 5*E-K*). Islets from 8 wk old SPCA1^+/-^ mice exhibited increased Phase 1 amplitude (Fig. 5*H*) and period (Fig. 5*I*) in response to glucose. During the Oscillatory Phase, SPCA1^+/-^ islets showed increased oscillation amplitude (Fig. 5*J*) and a decreased oscillation period (Fig. 5*K*) compared to WT islets.

### SPCA1 Deficiency Reduces Autophagy in β cells

Given the central role of the Golgi apparatus in insulin granule biogenesis, we next assessed insulin granule morphology in WT and SPCA1^+/-^ islets using electron microscopy (Fig. 6*A*). Islets from SPCA1^+/-^ mice exhibited a lower percentage of mature granules and a higher percentage of immature granules than WT islets (Fig. 6*B*). To help determine the mechanism behind impaired insulin secretion and altered granule maturation state, we performed a reverse phase protein array (RPPA) on WT and SPCA1KO INS-1 cells (Supplemental File 1). RPPA fold change results were used for pathway analysis with Metascape (Supplemental File 1). Pathway analysis revealed enrichment of several autophagy-related pathways, including “autophagy,” “mTOR signaling pathway,” “macroautophagy,” “selective autophagy,” and “mitophagy – animal” (Fig. 6*C*).

**Figure 6.**
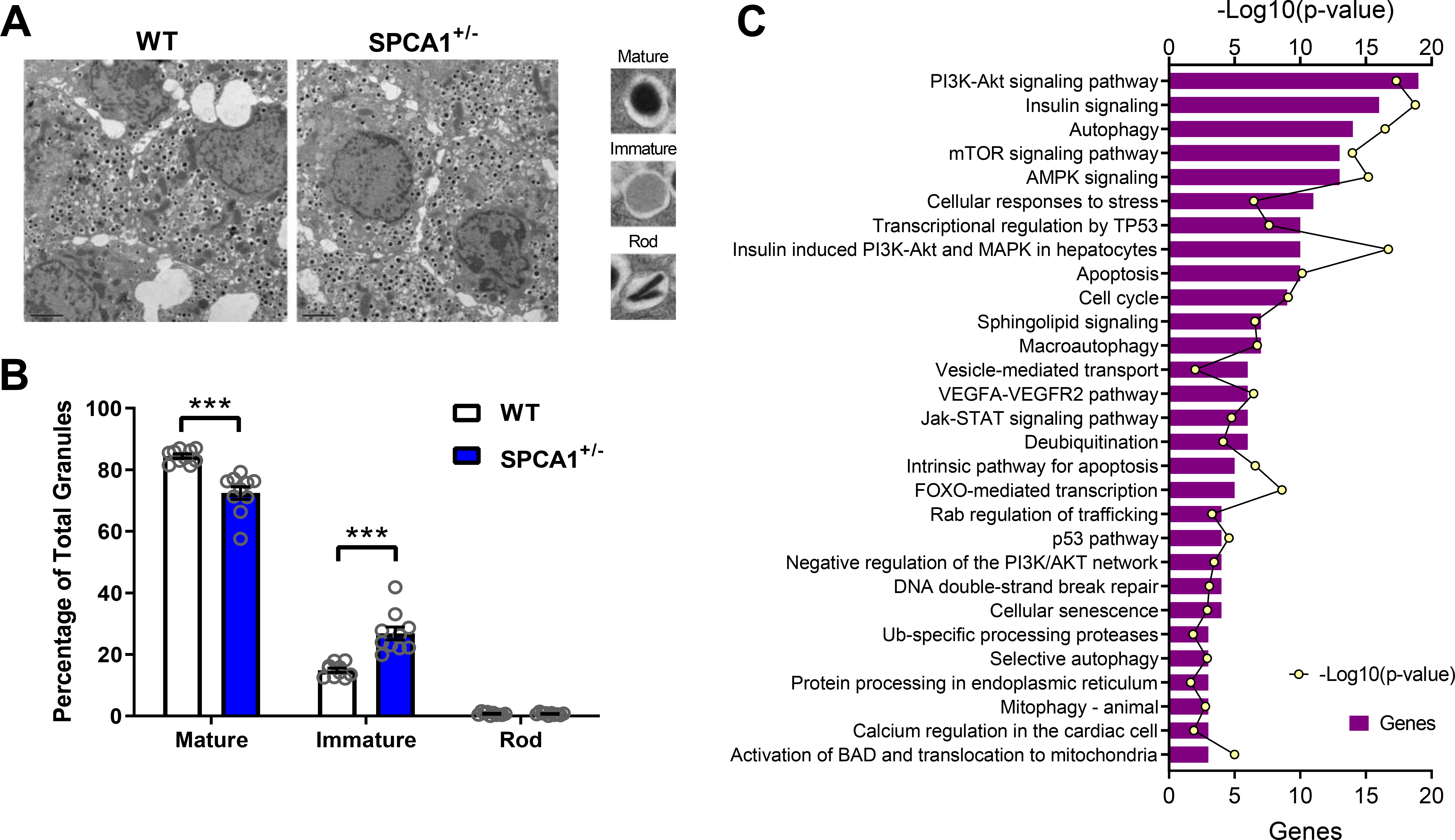
SPCA1 reduction alters insulin granule maturation state and autophagy-related pathways. (*A*-*B*) Islets from 8 wk old male WT and SPCA1^+/-^ mice were isolated and pooled for electron microscopy. Mature granules appear with a black or dark grey core surrounded by a halo. Immature granules have a grey core and lack a pronounced halo around the core. Rod-shaped granules are black and appear as rods within the vesicle. (*A*) Representative electron micrographs. Scale bar = 2 μm. (*B*) Quantification of mature, immature, and rod-shaped insulin granules. Islets were pooled from 2-3 mice and insulin granules counted in 10 fields of view per genotype, \*\*\**P* < 0.001, unpaired *t*-tests. (*C*) WT and SPCA1KO cells were processed for a Reverse Phase Protein Array (RPPA). RPPA results were processed with Metascape for pathway analysis. Representative pathways displayed with the corresponding number of significant genes (purple bar) and *P*-value (yellow dot).

Based on these results and existing reports demonstrating the regulatory role of autophagy in the turnover of insulin granules [35, 36], we tested whether autophagy pathways were modified by the loss of SPCA1. For this purpose, pancreas sections from 8 wk old WT and SPCA1^+/-^ mice were stained for insulin and the autophagy marker LC3 (Fig. 7*A*; Fig. S2). Fewer LC3 puncta were observed in SPCA1^+/-^ mouse islets than in WT littermates, suggesting reduced autophagy in haploinsufficient mice (Fig. 7*B*). Furthermore, SPCA1^+/-^ islets exhibited fewer insulin granules contained within LC3 puncta (Fig. 7*C*), possibly suggesting impaired vesicophagy [37].

**Figure 7.**
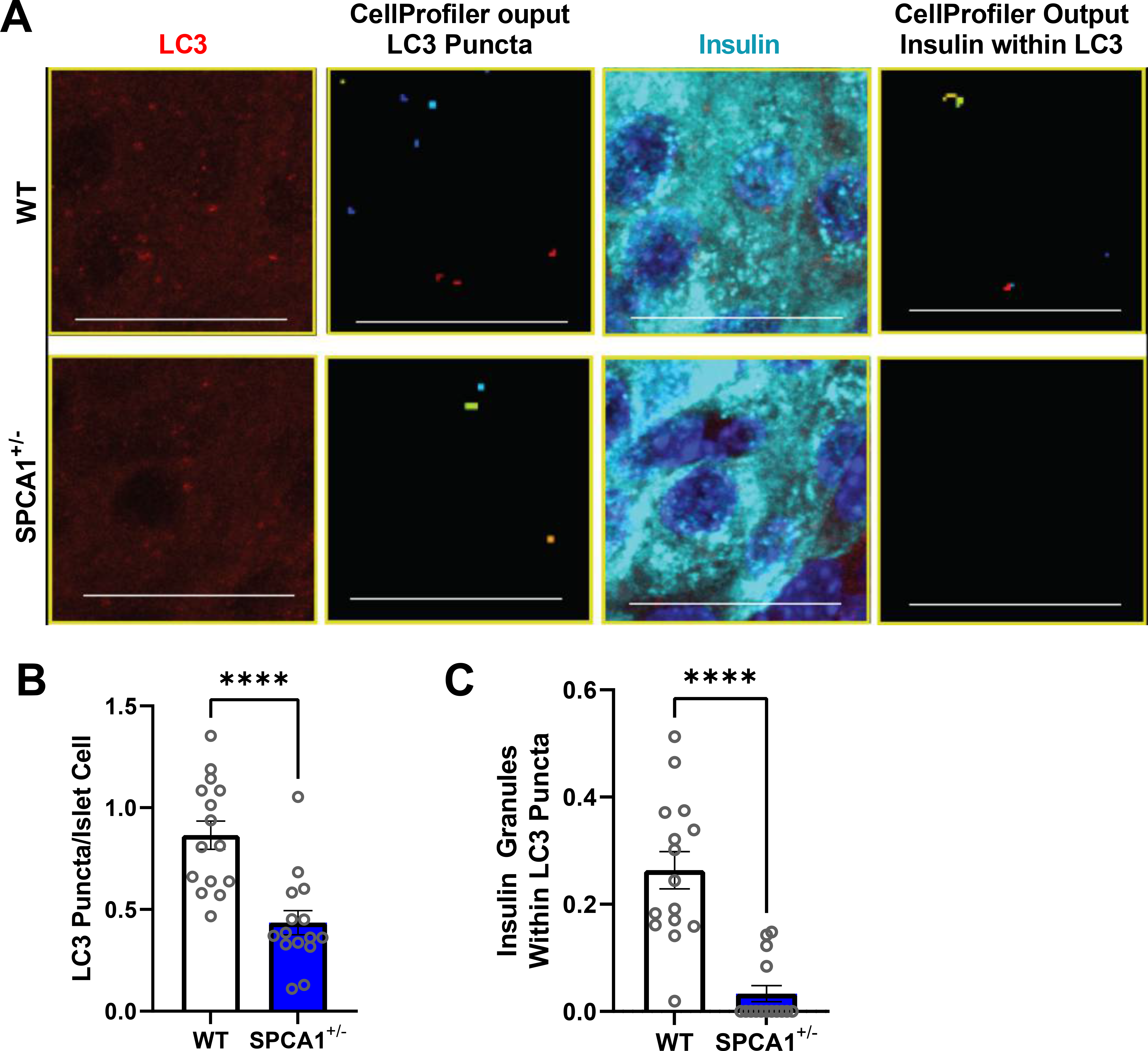
Autophagy is reduced in SPCA1^+/-^ β cells. (*A*) Pancreas sections from 8 wk old WT and SPCA1^+/-^ mice were stained for LC3 and insulin, and immunofluorescent images were acquired. Representative confocal images and CellProfiler output images are shown; scale bar = 20 μm. (*B*) LC3 puncta per islet area; *n* = 5 islets from 3 mice per genotype, \*\*\**P* < 0.001, unpaired *t*-test. (*C*) Insulin-containing LC3 puncta per islet; *n* = 5 islets from 3 mice per genotype, \*\*\*\**P* < 0.0001, unpaired *t*-test.

To further investigate autophagy in cells with reduced SPCA1, we activated autophagy in WT and SPCA1KO INS-1 cells by inhibiting the mTOR pathway with Torin1. SPCA1KO INS-1 cells had reduced expression of the autophagosome marker LC3-II (Fig. 8*A-B*) and a trend towards upregulated expression of p62, a marker of proteins tagged for autophagy-dependent degradation (Fig. 8*C-D*). Similarly, islets from SPCA1^+/-^ mice had a reduced level LC3-II following autophagy induction with Torin1 (Fig. 8*E-F*) and a trend towards upregulation of p62 (Fig. 8*G-H*).

**Figure 8.**
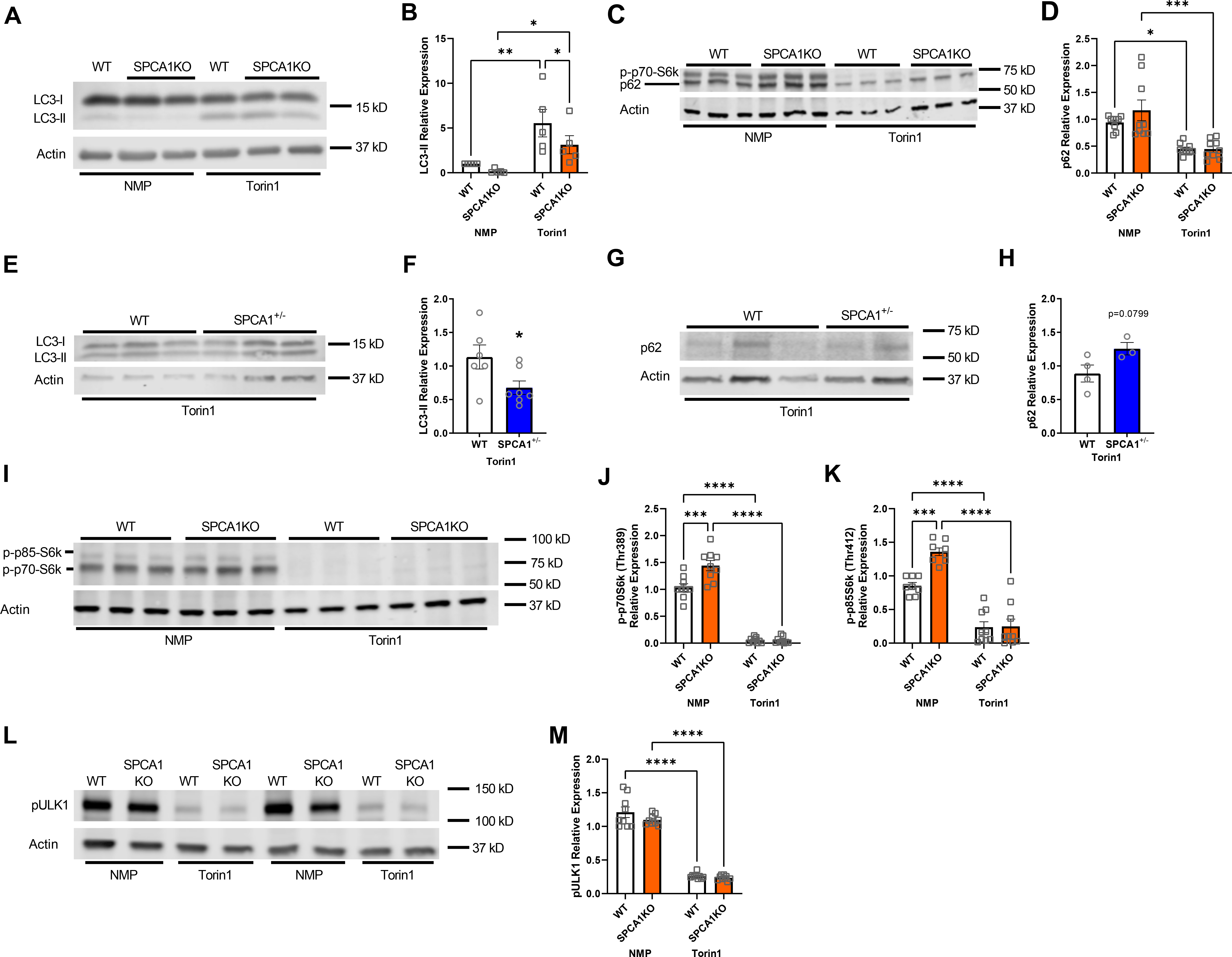
Induction of autophagy is impaired in SPCA1-deficient β cells. (*A-D*) WT and SPCA1KO INS-1 cells were treated with vehicle (NMP) or 250 nM Torin1 for 8 h. (*A*) Representative immunoblot showing LC3-II protein expression in NMP- and Torin1-treated cells. (*B*) Quantification of LC3-II protein expression, normalized to actin; *n* = 5, \**P* < 0.05, \*\**P* < 0.01, two-way ANOVA. (*C*) Representative immunoblot showing p62 protein expression in NMP- and Torin1-treated cells. (*D*) Quantification of p62 protein expression, normalized to actin; *n* = 9, \**P* < 0.05, \*\*\**P* < 0.001, two-way ANOVA. (*E-H*) WT and SPCA1^+/-^ islets were treated with 250 nM Torin1 for 8 h. (*E*) Representative immunoblot showing LC3-II protein expression in treated islets. (*F*) Quantification of LC3-II protein expression normalized to actin; *n* = 6-7; \**P* < 0.05, unpaired *t*-test. (*G*) Representative immunoblot showing p62 protein expression in treated islets. (*H*) Quantification of p62 protein expression, normalized to actin; *n* = 3-4. (*I-M*) WT and SPCA1KO INS-1 cells were treated with vehicle (NMP) or 250 nM Torin1 for 8 h. (*I*) Representative immunoblot showing p-p70-S6k and p-p85-S6k protein expression in NMP- and Torin1-treated cells. Quantification of p-p70-S6k (*J*) and p-p85-S6k (*K*) protein expression normalized to actin; *n* = 9; \*\*\**P* < 0.001, \*\*\*\**P* < 0.0001, two-way ANOVA. (*L*) Representative immunoblot showing pULK1 protein expression in NMP- and Torin1-treated cells. (*M*) Quantification of pULK1 protein expression, normalized to actin; *n* = 9; \*\*\*\**P* < 0.0001, two-way ANOVA.

Since the RPPA pathway analysis implicated an altered mTOR pathway in SPCA1KO and mTOR suppresses autophagy, we next assessed the phosphorylation of p70S6k and its isoform p85S6k downstream of mTOR. Both p-p70S6k and p-p85S6k were reduced in Torin1-treated SPCA1KO INS-1 cells (Fig. 8*I-K*), suggesting that the suppression of autophagy depended on the activation of the mTOR pathway. We also assessed the phosphorylation of ULK1, which is essential to autophagosome formation and is negatively regulated by mTOR. While not significant, a slight decrease in p-ULK1 occurred in vehicle-treated SPCA1KO INS-1 cells (Fig. 8*L-M*).

To further assess the effect of SPCA1 loss on autophagy flux, WT and SPCA1KO INS-1 cells were treated with 3-MA (Fig. S3*A-C*), a phosphoinositide 3-kinase inhibitor that prevents autophagy by blocking autophagosome formation, or with bafilomycin A and chloroquine (Fig. S3*D-F*), which prevent autophagosome-lysosome fusion and autophagosome cargo degradation. Following 3-MA treatment, LC3-II (Fig. S3*B*) and p62 (Fig. S3*C*) levels were comparable in WT and SPCA1KO INS-1 cells suggesting that SPCA1 does not affect the autophagosome formation. Similarly, LC3-II (Fig. S3*E*) and p62 (Fig. S3*F*) levels did not differ between WT and SPCA1KO INS-1 cells after bafilomycin A and chloroquine treatment, suggesting that SPCA1 is not implicated in autophagosome degradation.

## DISCUSSION

Calcium partitioning within the β cell is closely regulated at the level of the secretory pathway organelles. Within this compartment, Ca^2+^ regulates the egress of secreted proteins and acts as an important cofactor for chaperones, foldases, and a variety of enzymes essential for cargo processing and maturation [38, 39]. Secretory pathway Ca^2+^-dependent enzymes play a critical role in insulin production within the β cell. In the ER, proinsulin disulfide bonds are formed by protein-disulfide isomerase (PDI) [40]; and in the trans-Golgi and secretory granules, prohormone convertase 1/3 (PC1/3), PC2, and carboxypeptidase E (CPE) participate in insulin maturation [41, 42]. Intraluminal Golgi Ca^2+^ levels are maintained primarily by the Ca^2+^ pump SPCA, but the contribution of Golgi Ca^2+^ and SPCA to β cell function is largely unknown. Herein, we sought to address this knowledge gap by utilizing an INS-1 cell line lacking SPCA1 and a mouse model of SPCA1 haploinsufficiency.

In this study, we identified SPCA1 as the predominant isoform in the β cell and documented that SPCA1 is reduced in islets from multiple models of diabetes, suggesting a possible contribution to β cell function. Loss of SPCA1 resulted in reduced intraluminal Golgi Ca^2+^ in both the medial- and trans-Golgi compartments and increased baseline cytosolic Ca^2+^, consistent with previous reports [17,43,44]. Importantly, this reduction in Ca^2+^ levels was independent of Golgi volume reduction, further supporting the finding that SPCA1 loss reduced Golgi Ca^2+^ concentration. SPCA1 expression is highest in the trans-Golgi and lowest in the cis-Golgi, where SERCA is also expressed and partially contributes to Ca^2+^ level maintenance [11, 16]. Nonetheless, SPCA1 loss is sufficient to significantly lower Golgi Ca^2+^ levels.

Interestingly, earlier studies [17, 45] concluded that SPCA1 knockdown with siRNA increased GSIS, whereas our findings suggest that SPCA1KO INS-1 cells and SPCA1^+/-^ islets exhibit impaired GSIS. We believe this discrepancy reflects the difference between the effects of acute and chronic loss of SPCA1. The acute reduction in SPCA1 and Golgi Ca^2+^ caused by siRNA could be analogous to the secretory pathway Ca^2+^ release into the cytosol that amplifies insulin secretion [34, 46]. Conversely, our SPCA1KO INS-1 cells and haploinsufficient islets manifest a chronic reduction in SPCA1 and Golgi Ca^2+^ levels, resulting in the impaired GSIS we observed. This is similar to the effect observed in other secretory cell types, where SPCA1 loss has been associated with decreased secretion. For example, lactation requires high mammary gland Golgi Ca^2+^ wherein SPCA1 is upregulated during lactation and rapidly downregulated after lactation cessation [47, 48].

While we observed reduced GSIS in isolated haploinsufficient islets and SPCA1KO INS-1 cells, we did not observe overt defects in glucose tolerance or body composition in SPCA1^+/-^ mice. Even after subjecting WT and SPCA1^+/-^ mice to a high-fat diet, no changes in glucose tolerance, insulin tolerance, or body composition were observed (Fig. S1). These data suggest that SPCA1 haploinsufficiency is not adequate to impact whole-body glucose homeostasis. However, since diabetes development is multifactorial, and we observed reduced expression of SPCA1 in human islets from donors with T2D, loss of SPCA1 could contribute to reduced β cell function under diabetic conditions.

In addition to reduced insulin secretion, we observed increased insulin content at both the transcript and protein levels in INS-1 cells. While this increase was not phenocopied in the whole pancreas or isolated islets from SPCA1^+/-^ mice, insulin granule morphology was altered in SPCA1^+/-^ islets, which have a lower percentage of mature and a higher percentage of immature insulin granules present in electron micrographs. This observation led us to advance two possibilities: either insulin processing is impaired during maturation, or homeostasis of insulin granules is altered. We opted to explore the latter because the turnover of insulin granules has been proposed to be regulated by autophagy [35,36,49]. In further support of this notion, no difference was observed in the number of rod-shaped granules, which have been suggested to increase during impaired processing [21, 50]. In pancreas sections from SPCA1^+/-^ mice, we observed fewer LC3 puncta, which is consistent with LC3 undergoing lipidation and becoming part of the active autophagosome [51]. We also found several autophagy-related pathways altered by the loss of SPCA1. To further assess autophagy, we stimulated autophagy by inhibiting the mTOR pathway with Torin1 and observed reduced LC3-II in INS-1 cells and SPCA1^+/-^ islets. These results imply that cells lacking SPCA1 have impaired initiation of autophagy, a conclusion supported by the increase of phospho-p70S6k, a known suppressor of autophagy in the mTOR pathway, in SPCA1KO cells. Furthermore, when autophagosome formation or autophagosome-lysosome fusion and acidification were inhibited, we did not observe changes in LC3-II or p62, further suggesting that autophagy initiation is impaired by SPCA1 deficiency. Golgi Ca^2+^ may also regulate β cell autophagy; however, evidence of autophagy control by intracellular Ca^2+^ is conflicting [52, 53]. Several reports suggest that autophagy is upregulated by increased cytosolic Ca^2+^, via the CaMKKβ/AMPK signaling pathway [54-56]. Conversely, Ca^2+^ has been suggested to inhibit autophagy by calpain-mediated Atg5 proteolysis [57] or direct activation of mTORC1 [58]. Future investigations using SPCA1-deficient models with reduced intraluminal Golgi Ca^2+^ could help delineate this apparent discrepancy.

While SPCA1 deficiency has not been previously associated with impaired β cell function in humans, SPCA1 deficiency has been linked with a skin disease known as Hailey-Hailey disease. Mutations in one allele of *ATP2C1*, the gene encoding for SPCA1, lead to skin lesions caused by defects in the cell-to-cell adhesion and separation of epidermal layers [59]. Keratinocytes of individuals with Hailey-Hailey disease exhibit decreased Golgi Ca^2+^ levels and, like the β cell, rely on Golgi Ca^2+^ for essential cell functions, including secretion of proteins [59-61]. While multiple Hailey-Hailey case studies have listed diabetes as comorbidity [62-64], whether individuals with Hailey-Hailey disease exhibit increased susceptibility to diabetes has not been rigorously tested in large cohort studies.

In conclusion, our data reveal reduced expression of SPCA1 within the islet in multiple models of diabetes, including human islets from organ donors with T2D. We show for the first time that loss of SPCA1 leads to reduced intraluminal Golgi Ca^2+^ in β cells, reduced glucose-induced insulin secretion, and alterations in insulin granule maturation that were linked with altered mTOR pathways and impairments in the induction of autophagy.

## AUTHOR CONTRIBUTIONS

Designed research studies (RNB, TK, CEM), conducted experiments (RNB, XT, TK), acquired data (RNB, TK), analyzed data (RNB, SAW, CM, PK, TK, CEM), edited the manuscript (RNB, XT, SAW, CM, PK, TK, CEM), wrote the manuscript (RNB, CEM). CEM is the guarantor of this work and, as such, had full access to all the data in the study and takes responsibility for the integrity of the data and the accuracy of the data analysis.

## ACKNOWLEDGEMENTS

The authors would like to thank Solaema Taleb, MS, for technical assistance, Gary E. Shull, PhD for reagents and reading the manuscript, Richard N. Day, PhD, for assistance with fluorescence lifetime imaging microscopy studies, and Timothy A. Reinhardt, PhD, for reagents. The authors would like to thank the Indiana Diabetes Research Center Islet & Physiology Core (P30 DK097512) for data obtained using core equipment and services. This work was supported by the Histology Core of the Indiana Center for Musculoskeletal Health at IU School of Medicine. This work was supported by the National Institute of Diabetes and Digestive and Kidney Diseases (NIDDK) grants R01 DK093954 and UC4 DK104166 (to CEM), U.S. Department of Veterans Affairs Merit Award I01BX001733 (to CEM), and gifts from the Sigma Beta Sorority, the Ball Brothers Foundation, and the George and Frances Ball Foundation (to CEM). RNB was supported by a National Institute of Allergy and Infectious Diseases training grant (T32 AI060519) and by a JDRF postdoctoral fellowship (3-PDF-2017-385-A-N). Human pancreatic islets were provided by the NIDDK-funded Integrated Islet Distribution Program (IIDP) (RRID:SCR_014387) at City of Hope, NIH grant (2UC4 DK098085).

## PRIOR PRESENTATION

Parts of this study were presented at the 77^th^ American Diabetes Association Scientific Sessions June 9-13, 2017, the 78^th^ American Diabetes Association Scientific Sessions June 22-26, 2018, the HIRN 2017 Annual Investigator Meeting March 7-10, 2017, the 2016 Midwest Islet Club Conference May 18-19, 2016, and the 2021 Midwest Islet Club Conference July 13-15, 2021.

## SUPPORTING INFORMATION

### Animal studies

For the high-fat diet study (Fig. S1), mice were fed a high-fat diet containing 45% kilocalories from fat or a sucrose-matched control diet containing 10% kilocalories from fat (Research Diets, Inc., New Brunswick, NJ) beginning at 8 wks of age until 24 wks of age.

### Immunofluorescence, CellProfiler analysis, and β cell area

For immunofluorescence of cultured cells, cells were fixed with 4% paraformaldehyde (PFA), permeabilized with 0.25% Triton X-100, and blocked with animal-free blocking buffer (Vector Laboratories, Burlingame, CA). For immunofluorescence of the pancreas, the tissue was PFA-fixed paraffin-embedded, and 6 μm sections were cut. The sections were deparaffinized, subjected to antigen retrieval using citrate-based Antigen Unmasking Solution (Vector Laboratories), and blocked with animal-free blocking buffer. Cells or pancreas sections were then incubated overnight at 4°C with primary antibodies (Table S2), followed by Alexa fluor-conjugated secondary antibodies. Nuclei were stained with DAPI. Z-stack images were obtained with a Zeiss confocal microscope. For Golgi volumetric analysis, z-stack images were imported into ImageJ, stacks were summated with the “sum slices” function, the threshold was equally adjusted with the “max entropy” function, and the area was recorded and normalized to the number of nuclei in the image.

For LC3 analysis, fluorescence intensity measurements, puncta counts, and colocalization analyses were performed using CellProfiler 4.1.3 (cellprofiler.org) [1]. Background was subtracted from each image by removing the lower-quartile intensity from each channel. Pancreas regions of interest (ROIs) were defined by insulin-positive area, and LC3 puncta were identified by discarding objects outside a pixel diameter range after median filtering [2]. The puncta count was normalized to the insulin-positive area. For colocalization positivity, parent (insulin) and child (LC3) objects were defined [2] and the percentage of LC3 puncta colocalized or co-compartmentalizing with insulin granules was determined using CellProfiler’s built in tutorials.

For the β cell area, pancreatic sections were prepared as above and incubated with anti-insulin primary antibody overnight at 4°C. Insulin was visualized with anti-rabbit ImmPRESS reagent and NovaRed substrate kit; hematoxylin was used to counterstain tissue. Brightfield images were acquired with a Zeiss AxioScan.Z1 microscope (Zeiss, Oberkochen, Germany).

### Live Cell Imaging

For the Calcium 6 assays, 10,000 cells/well were seeded in black wall, clear bottom 96-well plates (Costar, Tewksburry, MA) and cultured for two days. Calcium 6 reagent was added to cells, and after incubation for 2 h at 37°C, the medium was replaced with Ca^2+^-free HBSS supplemented with 0.2% BSA and EGTA. The assay was performed on the FlexStation 3 system at 37°C/5% CO_2_ using an excitation wavelength of 485 nm and an emission wavelength of 525 nm. The baseline was recorded for 30 s at a 1.52 s reading interval; subsequently, drugs were added and the recording continued until 200 s. For data analysis, the ΔF/F0 was derived from the drug response (ΔF) divided by the resting intracellular Ca^2+^ (F0).

For fluorescence lifetime imaging microscopy (FLIM), cells (3 x 10^6^/well) were seeded in Lab-Tek II Chambered Coverglass (Nunc, Roskilde, Denmark) and transduced with 8 x 10^6^ viral particles/mL adenovirus for 24 h prior to imaging. Golgi compartment-specific adenoviruses expressing Cameleon calcium sensor probes specific for the medial-Golgi, driven by the 1,6 N acetyl-glucosaminyl transferase promoter [3] and the trans-Golgi, driven by the sialyl-transferase1 promoter [4], were generated by the Human Islet and Adenovirus Core at Einstein-Mount Sinai Diabetes Research Center (Bronx, NY). The Alba FastFLIM system (ISS Inc., Champaign, IL) was coupled to an Olympus IX71 with a 60x water-immersion lens (Olympus, Tokyo, Japan). Confocal imaging at 530/43 nm acceptor and 480/40 nm receptor wavelengths was performed with VistaVision software (ISS Inc.). ROIs were selected with >100 count averages and lifetimes were obtained by analyzing the first 8-10 modulation frequencies. The efficiency of FRET was estimated using the following equation: *E*_FRET_ = 1 - (τ_DA_ / τ_D_), where τ_D_ and τ_DA_ are the donor fluorescence lifetime obtained in the absence and presence of the acceptor, respectively.

### Buffer and Media Formulations

The INS-1 medium was composed of RPMI-1640 (containing 11.1 mM glucose), 10% FBS, 100 U/0.1 mg/L penicillin/streptomycin, 10 mM HEPES, 2 mM L-glutamine, 1 mM sodium pyruvate, and 50 μM β-mercaptoethanol.

The islet medium was composed of phenol red-free DMEM (containing 5.5 mM glucose), 10% FBS, 10 mM HEPES, 2 mM L-glutamine, and 100 U/0.1 mg/L penicillin/streptomycin.

Protein Lysis Buffer consisted of 0.05% deoxycholate, 0.5% NP40, 0.1% SDS, 0.2% Sarcosyl, 10% glycerol, 1 mM DTT, 1 mM EDTA, 10 mM NaF, 10 mM Tris-pH 8.0, and volumed with ultrapure H_2_O. 10% SDS, 10% PhosSTOP, 10% cOmplete protease inhibitor, and 0.1% Benzonase were added immediately prior to use.

Protein Loading Buffer was made from an equal mixture of 6x sample buffer (47% glycerol, 416 μM SDS, 0.9 nM bromophenol blue, 60 mM Tris-pH 6.8, and volumed with ultrapure H_2_O) and 2x Laemmli sample buffer (Biorad #1610737) for a final 4x working solution. Immediately before use, 20 μL of 1 M DTT and 30 μL β-mercaptoethanol was added to 1 mL of 4x working solution.

RPPA Lysis Buffer consisted of 1% Triton X-100, 50 mM HEPES, 150 mM NaCl, 1.5 mM MgCl_2_, 1 mM EGTA, 100 mM NaF, 10 mM Na-pyrophosphate, 1 mM Na_3_VO_4_, 10% glycerol, pH 7.4 and was volumed with ultrapure H_2_O. 10% PhosSTOP and 10% cOmplete protease inhibitor were added immediately before use.

RPPA 4x Sample Buffer was composed of 40% glycerol, 8% SDS, 0.25 M Tris-HCl, pH 6.8, and 10% 2-mercaptoethanol.

Electron Microscopy Fixative contained 2% glutaraldehyde and 4% paraformaldehyde in 0.1 M sodium cacodylate buffer, pH 7.4.

Islet Ca^2+^ Imaging HBSS was comprised of HBSS supplemented with 2% BSA, 10 mM HEPES, 2 mM CaCl_2_, and 0.8 mM MgCl_2_.

Krebs buffer consisted of 114 mM NaCl, 4.7 mM KCl, 0.12 mM KH_2_PO_4_, 1.16 mM MgSO_4_, 20 mM HEPES, 2.5 mM CaCl_2_, 0.2% BSA, 25.5 mM NaHCO_3_, pH 7.2, and volumed with ultrapure H_2_O.

---

**Table S1.**
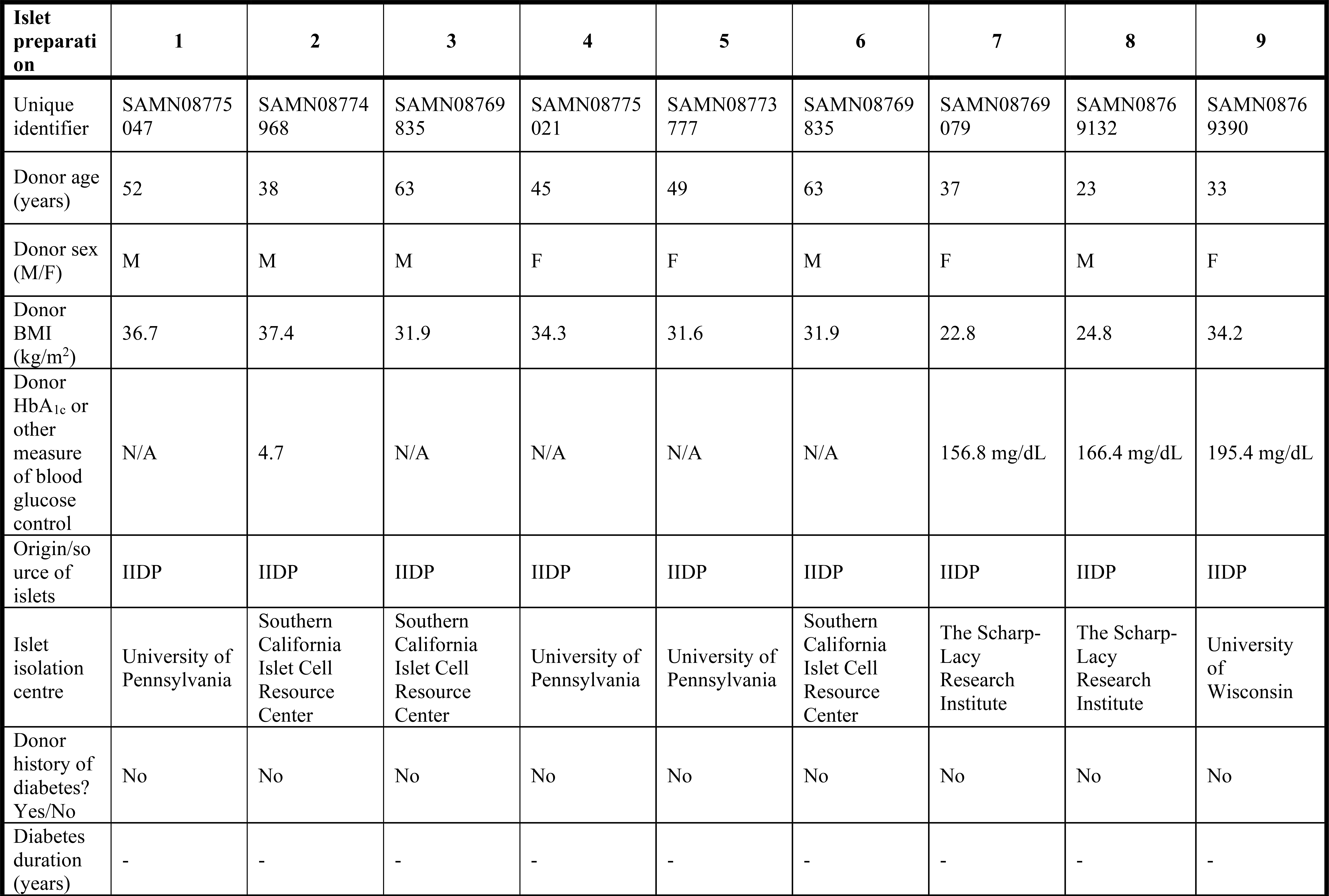
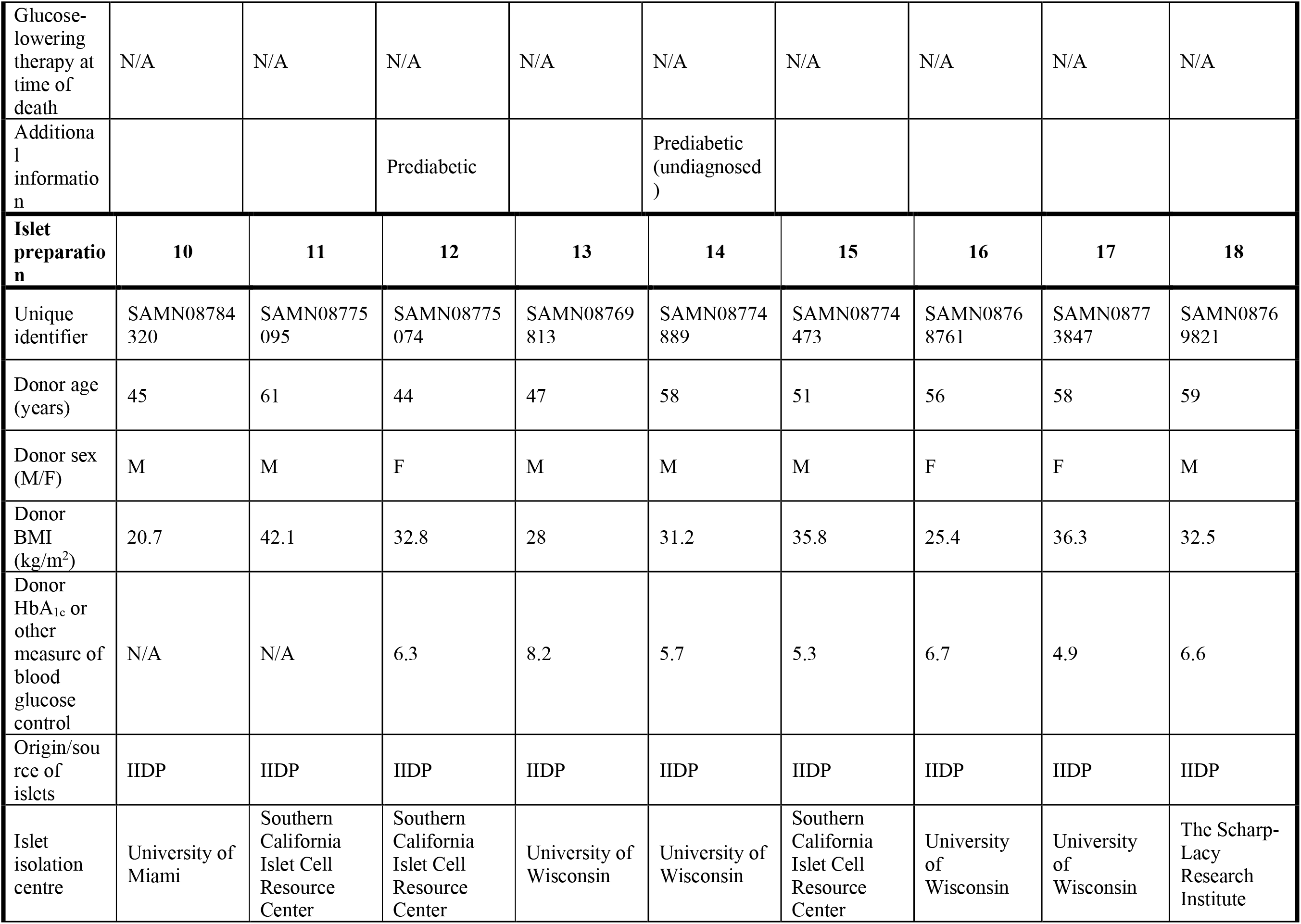
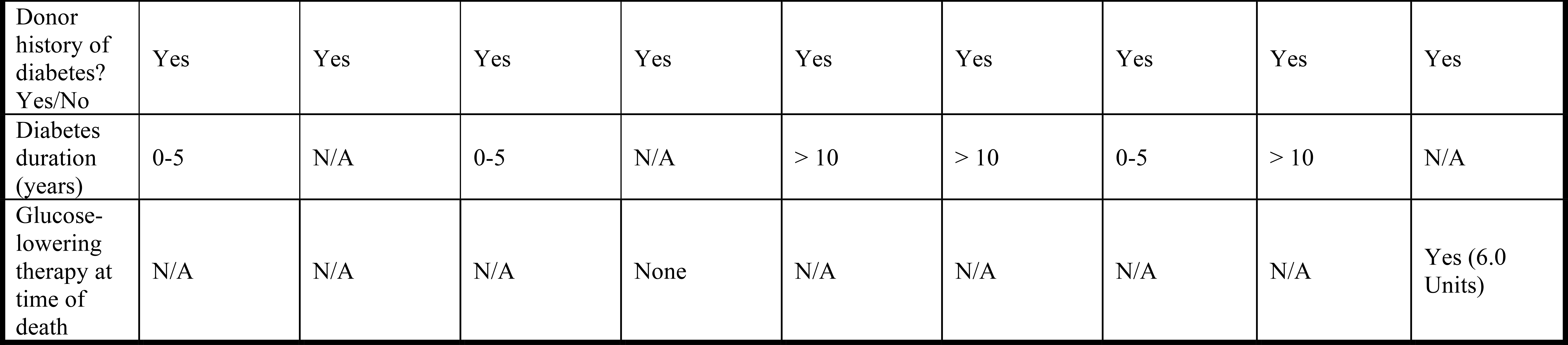
Human Cadaveric Islet Donor Characteristics.

**Table S2.**
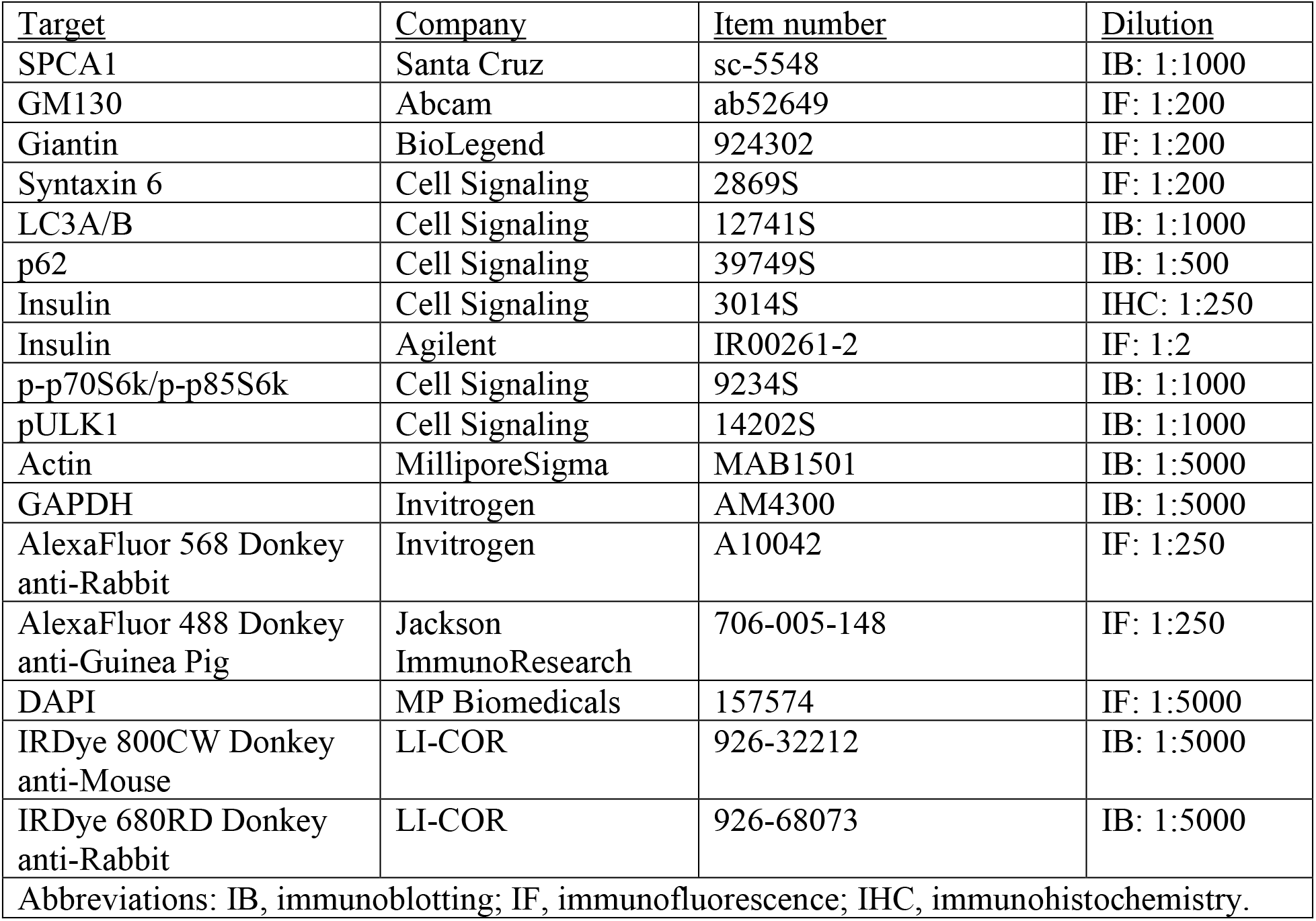
Antibodies.

| <u>Target</u> | <u>Company</u> | <u>Item number</u> | <u>Dilution</u> |
| --- | --- | --- | --- |
| SPCA1 | Santa Cruz | sc-5548 | IB: 1:1000 |
| GM130 | Abcam | ab52649 | IF: 1:200 |
| Giantin | BioLegend | 924302 | IF: 1:200 |
| Syntaxin 6 | Cell Signaling | 2869S | IF: 1:200 |
| LC3A/B | Cell Signaling | 12741S | IB: 1:1000 |
| p62 | Cell Signaling | 39749S | IB: 1:500 |
| Insulin | Cell Signaling | 3014S | IHC: 1:250 |
| Insulin | Agilent | IR00261-2 | IF: 1:2 |
| p-p70S6k/p-p85S6k | Cell Signaling | 9234S | IB: 1:1000 |
| pULK1 | Cell Signaling | 14202S | IB: 1:1000 |
| Actin | MilliporeSigma | MAB1501 | IB: 1:5000 |
| GAPDH | Invitrogen | AM4300 | IB: 1:5000 |
| AlexaFluor 568 Donkey anti-Rabbit | Invitrogen | A10042 | IF: 1:250 |
| AlexaFluor 488 Donkey anti-Guinea Pig | Jackson ImmunoResearch | 706-005-148 | IF: 1:250 |
| DAPI | MP Biomedicals | 157574 | IF: 1:5000 |
| IRDye 800CW Donkey anti-Mouse | LI-COR | 926-32212 | IB: 1:5000 |
| IRDye 680RD Donkey anti-Rabbit | LI-COR | 926-68073 | IB: 1:5000 |
| Abbreviations: IB, immunoblotting; IF, immunofluorescence; IHC, immunohistochemistry. |  |  |  |

**Table S3.**
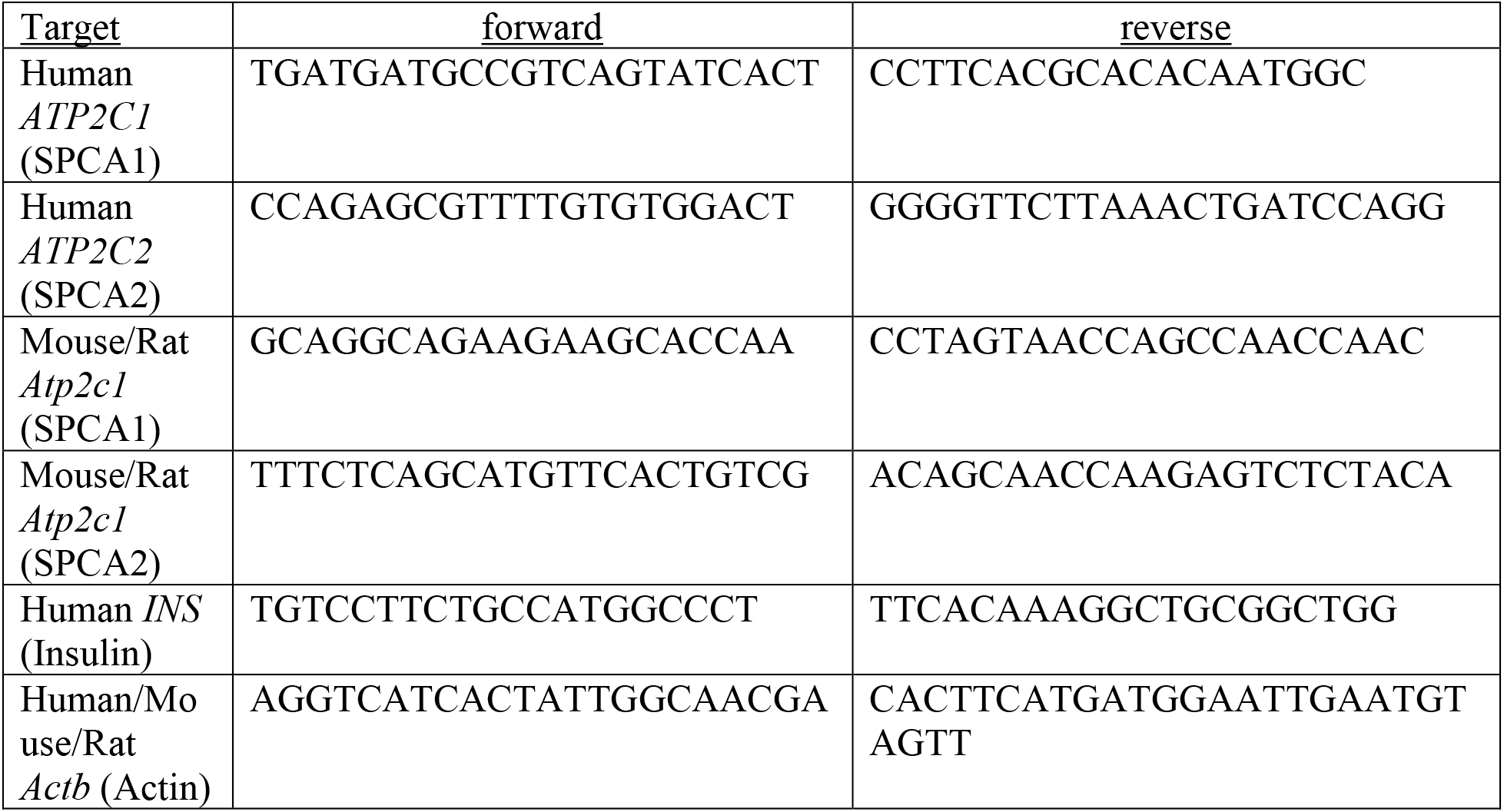
Primers.

**Table S4.**
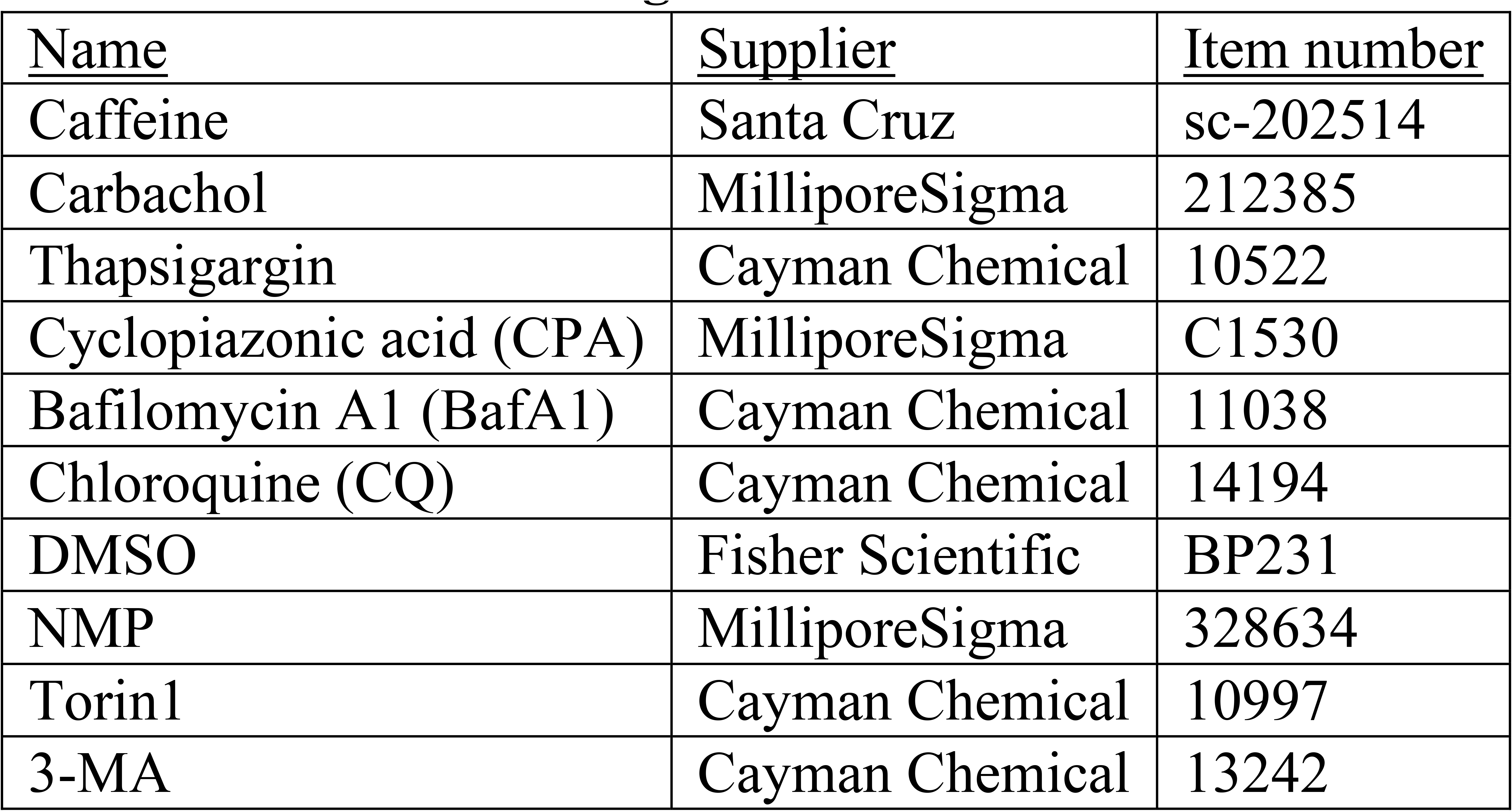
Treatment Reagents.

**Figure S1.**
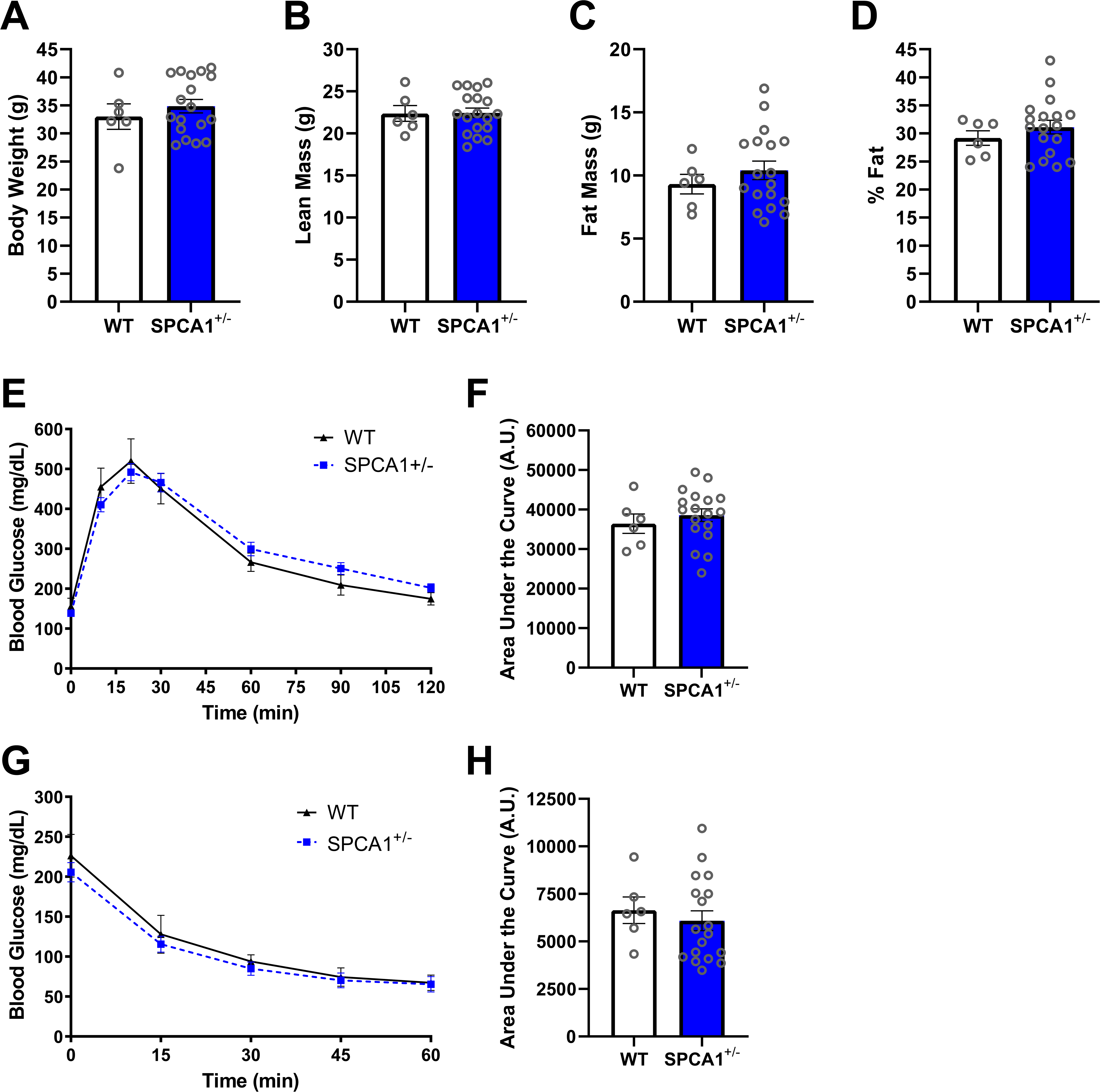
High-fat diet effects on body composition, glucose tolerance, and insulin tolerance. At 8 wks of age, WT and SPCA1^+/-^ mice were placed for 16 wks on high-fat diet (HFD) providing 45% kcal from fat. (*A*-*D*) EchoMRI body composition analysis of 24 wk old male WT and SPCA1^+/-^ mice after 16 wks of HFD. (*A*) Body weight; *n* = 6-18. (*B*) Lean mass; *n*=6-18. (*C*) Fat mass; *n* = 6-18. (*D*) Percent fat; *n* = 6-18. (*E*) Glucose tolerance test in 24 wk old male WT and SPCA1^+/-^ mice after 16 wks of HFD and (*F*) corresponding area under the curve (AUC); *n* = 6-18. (*G*) Insulin tolerance test in 23 wk old male WT and SPCA1^+/-^ mice after 15 wks of HFD and (*H*) corresponding area under the curve (AUC); *n* = 6-18.

**Figure S2.**
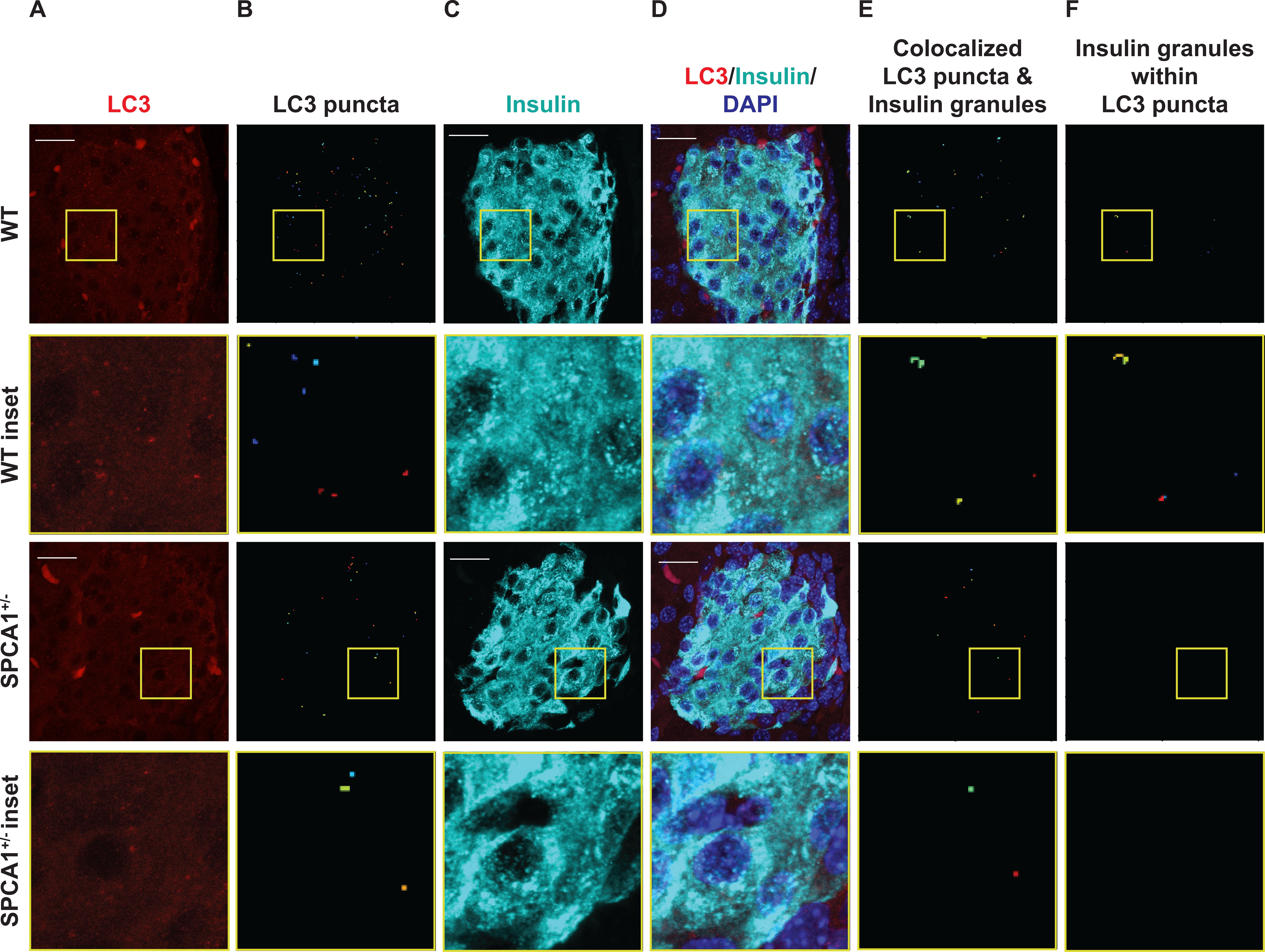
Representative immunofluorescent and CellProfiler images. (*A*-*F*) Pancreas sections from 8 wk old WT and SPCA1^+/-^ mice were stained for LC3 and insulin, and immunofluorescent images were acquired. Confocal images were processed with CellProfiler to identify LC3 puncta, insulin granules, LC3 puncta colocalized with insulin granules, and insulin granules surrounded by LC3 puncta. Confocal and CellProfiler output images show LC3 (*A*, confocal), LC3 puncta (*B*, CellProfiler output), insulin (*C*, confocal), merged LC3, insulin, and nuclei (*D*, confocal), LC3 puncta colocalized with insulin granules (*E*, CellProfiler output), and insulin granules surrounded by LC3 puncta (*F*, CellProfiler output). Scale bar = 20 μm.

**Figure S3.**
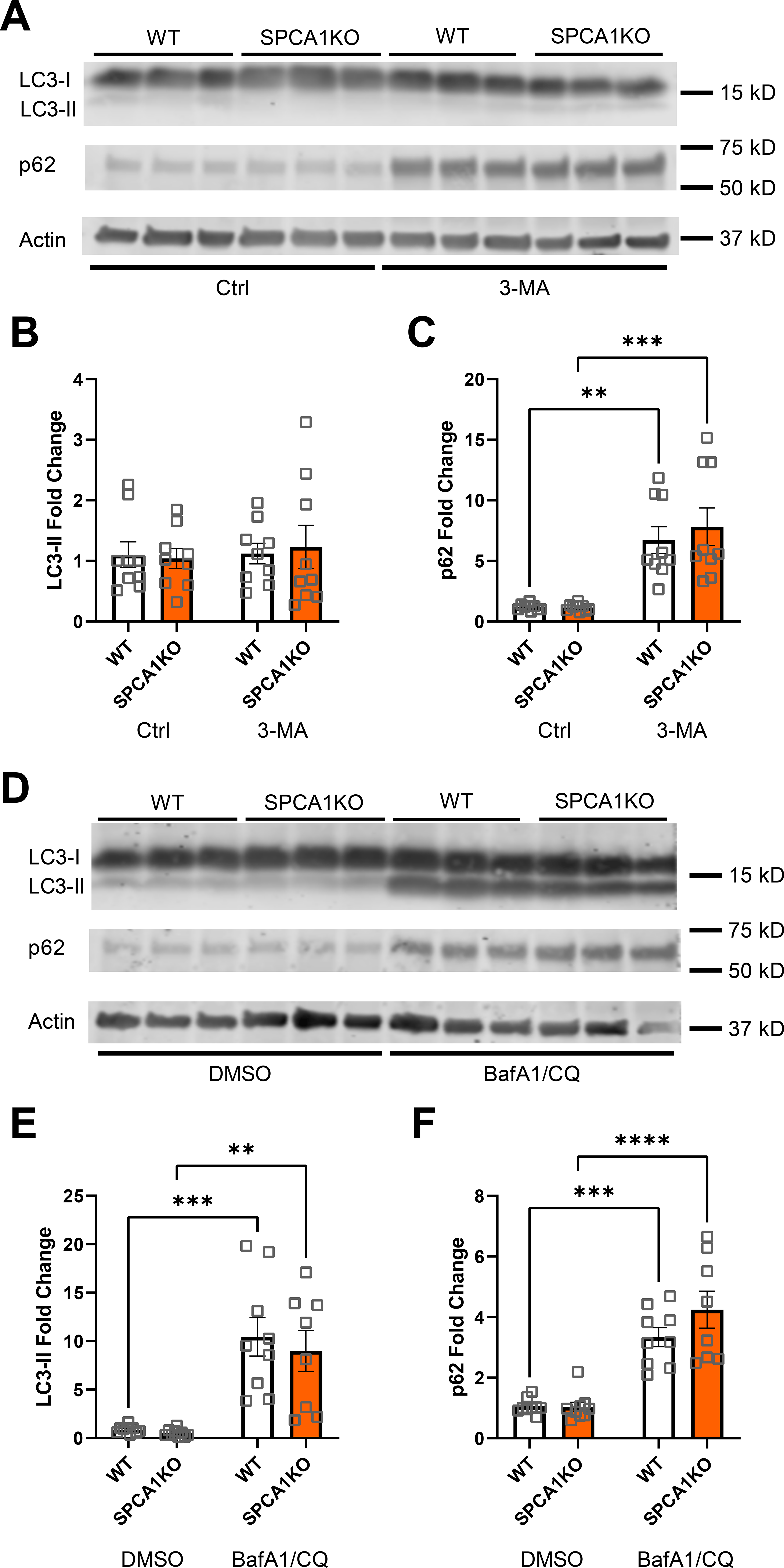
Autophagy after 3-MA or bafilomycin and chloroquine treatment is not altered by SPCA1 loss. (*A*-*C*) WT and SPCA1KO INS-1 cells were treated with media alone (Ctrl) or 5 mM 3-MA for 24 h. (*A*) Representative immunoblot showing LC3-II and p62 protein expression in control and treated INS-1 cells. Quantification of LC3-II (*B*) and p62 (*C*) protein expression, normalized to actin; *n* = 9, \*\**P* < 0.01, \*\*\**P* < 0.001, two-way ANOVA. (*D*-*F*) WT and SPCA1KO INS-1 cells were treated with vehicle (DMSO) or 10 nM bafilomycin A1 (BafA1) and 10 μM chloroquine (CQ) for 24 h. (*D*) Representative immunoblot showing LC3-II and p62 protein expression in control and treated INS-1 cells. Quantification of LC3-II (*E*) and p62 (*F*) protein expression, normalized to actin; *n* = 9; \*\**P* < 0.01, \*\*\**P* < 0.001, \*\*\*\**P* < 0.0001, two-way ANOVA.

